# Regulation of pupil size in natural vision across the human lifespan

**DOI:** 10.1101/2023.10.16.562484

**Authors:** Rafael Lazar, Josefine Degen, Ann-Sophie Fiechter, Aurora Monticelli, Manuel Spitschan

**Author notes:** To whom correspondence should be addressed: Prof. Dr Manuel Spitschan.

## Abstract

Vision is mediated by light passing through the aperture of the eye, the pupil, which changes in diameter from ~2 to ~8 mm from the brightest to the darkest illumination. In addition, with age, mean pupil size declines. In laboratory experiments, factors affecting pupil size can be experimentally controlled or held constant, but how the pupil reflects changes in retinal input from the visual environment under natural viewing conditions is not clear. Here, we address this question in a field experiment (*N*=83, 43 female, age: 18-87 years) using a custom-made wearable video-based eye tracker with a spectroradiometer measuring spectral irradiance in the approximate corneal plane. Participants moved in and between indoor and outdoor environments varying in spectrum and engaged in a range of everyday tasks. Our real-world data confirm that light-adapted pupil size is determined by light intensity, with clear superiority of melanopic over photopic units, and that it decreases with increasing age, yielding steeper slopes at lower light levels. We find no indication that sex, iris colour or reported caffeine consumption affect pupil size. Taken together, the data provide strong evidence for considering age in personalised lighting solutions and for using melanopsin-weighted light measures to assess real-world lighting conditions.

## Introduction

Human vision depends on light impinging on the retina and exciting the retinal photoreceptors. Serving as the aperture of the visual system, the pupil is able to adjust the amount of incident light by constricting or dilating to diameters ranging from ~2 to ~8 mm under the brightest and darkest conditions, respectively [1, 2]. By this route, the pupil regulates retinal illuminance within a limited range (~16x, or 1.2 log units between smallest and largest pupil) [3]. The pupil is also critical in attaining optimal optical quality of the retinal image [4, 5] by balancing necessary retinal illuminance with optical aberrations [6, 7] and depth of focus [8]. Hence, pupil size is an essential determinant for visual performance under changing ambient illumination.

### Light-level dependence of pupil size

It is well known that pupil size generally depends on light level. This effect is encoded by the joint action of the five retinal photoreceptor classes in the human retina: rods, three types of cones, and melanopsin, which is expressed in the intrinsically photosensitive retinal ganglion cells (*ipRGCs*). Each photoreceptor class is characterised by a distinct wavelength of maximal sensitivity, but due to their broad spectral tuning, the photoreceptors overlap greatly in their spectral sensitivities [9]. Rods have their peak sensitivity near 495 nm and characteristically show saturation under photopic or “daylight” light levels and are thus mainly responsible for scotopic or “night” vision. The three types of cones controlling photopic vision, namely the long-wavelength-sensitive L-cones, the medium-wavelength-sensitive M-cones and the short-wavelength-sensitive S-cones, peak near 558, 530, and 420 nm, respectively. Melanopsin exhibits its maximum spectral sensitivity near 480 nm. The ipRGCs expressing melanopsin differ markedly from the other photoreceptors in their temporal response to light, showing longer latency and sustained depolarization [10, 11]. While the ipRGCs are themselves photosensitive, they also receive synaptic inputs from the cones and rods [10]. All photoreceptors contribute to the control of the pupil, though their spatial distribution and temporal “niches” are different. Converging evidence suggests that in daylight light levels, the mean steady-state pupil size is mainly controlled by the melanopsin-encoded signal [11–16]. For dynamic pupil size responses, the melanopsin-encoded signal seems to have a smaller role in pupil size control with increasing temporal frequency [17, 18].

### Age-dependence of pupil size

Pupil size is affected by age. Steady-state pupil size starts to decrease with age after the second decade of life, both in dark [19–21] and light adaptation [13, 22–28]. This age effect, also termed “senile miosis”, is more pronounced in dimmer light compared to more intense light conditions [13, 24, 28, 29]. For instance, given a luminance of 9 cd/m², Winn et al. (1994) [28] reported an age related decrease of pupil size of ~0.043 mm per year. This amounts to a ~1.72 mm difference comparing a typical 60-year-old and 20-year old person at that light level. A variety of processes have been proposed as possible causes for the age-related functional decrease of the pupil [2] such as structural iris damage like stiffness and muscle degeneration [30, 31], sympathetic and parasympathetic deficits [23] as well as deficiency in central inhibition [21]. Notably, there are also various retinal degradation processes associated with ageing such as the decrease in photoreceptor density [32] and delayed photopigment regeneration [33], along with ageing phenomena like lens yellowing [34] that could partly contribute to the functional shrinking of pupil size.

### Pupil size model

Drawing from the extensive literature on stimuli affecting pupil size, Watson and Yellott (2012) [35] developed a unified formula for predicting pupil size from luminance, age, number of eyes and field diameter (see Fig. 1C for example predictions as a function of luminance and age). This formula summarises the luminance-dependence of the pupil from a total of eight extant studies [28, 36–42]. However, luminance reflects only a weighted combination of the L- and M-cones. As discussed previously, all photoreceptors participate in controlling pupil size and therefore, parametrizing retinal intensity as a function of luminance does not predict pupil size well [12, 15, 43].

**Figure 1:**
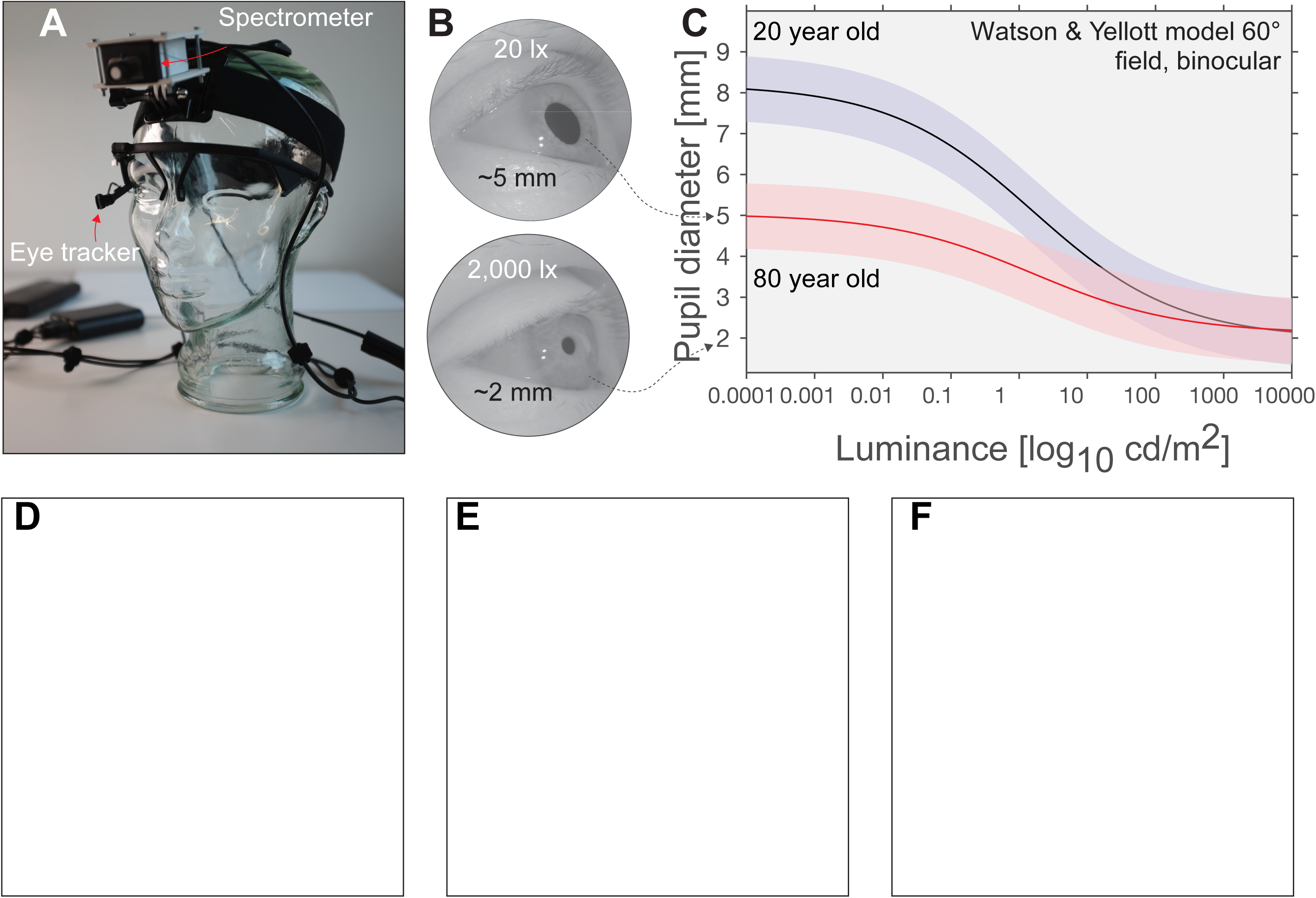
***A*** Glass mannequin head with ambulatory measurement setup, incorporating infra-red eye tracker, small-footprint spectrometer, as well as a control computer (Raspberry Pi) and a power bank. ***B*** Example images under two different illumination conditions varying across 3 orders of magnitude recorded with our setup. ***C*** Predictions of the [35] for two observers (20-year-old vs. 80-year-old). ***D-F*** Indoor (***D***), outdoor (***E***) and laboratory lighting situations (***F***).

### Pupil size in naturalistic conditions

Our understanding of the factors influencing pupil size almost exclusively stems from controlled laboratory studies using simplified parametric stimuli and has generated an internally valid body of well-controlled studies. Yet, to predict pupil size in the real world, research needs to be extended to naturalistic environments [44–46]. As pupil size is a crucial factor for visual performance in real life, it is necessary to confirm how it varies in an age- and gender-balanced sample, performing ecologically relevant tasks in everyday-life settings under naturalistic illumination.

To our knowledge, a very limited number of studies have yet investigated pupil size under real-world conditions. A study by Koch et al. (1991) [25] simulated viewing conditions of five naturalistic tasks in an indoor laboratory setting (night-time driving, reading dim and bright illumination, distant viewing in direct and indirect sunlight) and confirmed the decrease of pupil size with increased illumination and higher age. Harley and Sliney (2018) [47] examined outdoor daylight-adapted pupil sizes of 87 subjects from a military population between 790 cd/m^2^ and 4,250 cd/m^2^. Crucially, both studies parametrise the lighting conditions in terms of luminance (weighted sum of L- and M-cones). If a spectrum does not change with light level, then the change in (il)luminance as a function of light level will be proportional to the change in melanopic (ir)radiance. However, in the real world, the illuminant and reflectance spectra (and the resultant scene spectra) can be very diverse (the “spectral diet”, [48]), due to different sources of illumination (e.g. daylight, incandescent, fluorescent, LED lighting). As a consequence, (il)luminance and melanopic (ir)radiance are not always correlated (see Fig. 2 in [15] for this dissociation under real-world spectra).

**Figure 2:**
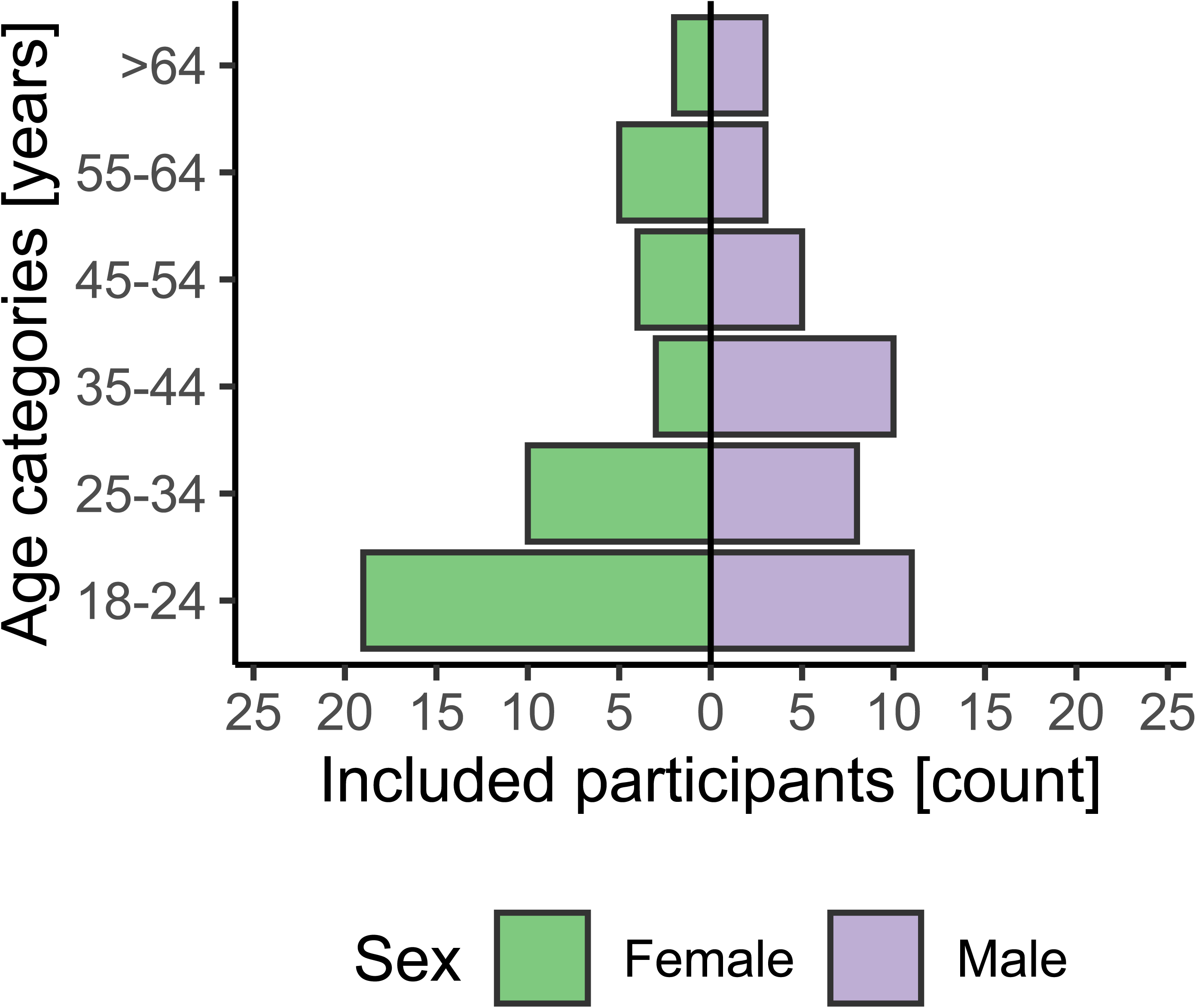
Included sample (n=83) stratified for age groups and sex. Green bars on the left depict female, and violet bars on the right depict male participants. The sample is skewed towards younger age groups, especially for female participants.

Here, we perform a confirmatory analysis of age effects and effects of spectrally weighted illumination on the light-adapted pupil size during real-life indoor and outdoor light and viewing-task conditions in a healthy, age-diverse sample. Our confirmatory hypotheses (CH) are:

- CH1: **In real-world conditions, light-adapted pupil size changes as a function of melanopsin sensitivity-weighted retinal illumination, with higher illumination associated with a smaller pupil size.**
- CH2: **In real-world conditions, melanopsin sensitivity-weighted retinal illumination better predicts pupil size than the weighted sum of L- and M-cone-weighted retinal illumination (CIE 1931 Y, photopic illuminance).**
- CH3: **In real-world conditions, light-adapted pupil size changes as a function of age, with higher age associated with a smaller pupil size.**

We tested each of these hypotheses using a linear mixed model (LMM) and calculated the total evidence for each of the hypotheses using the Bayes Factor (BF) comparing the full model formulated for each hypothesis with the appropriate null model (see *Analytic Strategy* below).

## Materials and Methods

### Participant characteristics

We aimed to recruit a total of 96 healthy volunteers (48 female, 48 male) from the community. To achieve a balanced and age-diverse sample, we sought to recruit 16 volunteers in each of the following age groups^1^: 18-24 years, 25-34 years, 35-44 years, 45-54 years, 55-65 years, and >65 years of age. Only participants wearing contact lenses or requiring no vision correction were included as our measurement device is not compatible with wearing glasses. All participants must have 20/40 visual acuity or better (assessed by Snellen chart) and normal colour vision (assessed by HRR test). Additionally, participants were screened for relevant medication, disorders (neurological, metabolic), eye conditions, mental health (assessed by the German version of the GHQ-12, [49]) and tested for consumption of drugs (urine-based multi-panel drug test, nal von minden; Moers, Germany) and alcohol (saliva-based alcohol test, ultimed; Ahrensburg, Germany). Elderly participants with intraocular lens replacements (IOLs) were allowed to be included in the study, with their lenticular status (i.e. whether they have an IOL) recorded as a binary variable^2^.

### Study protocol

Participants performed a range of natural tasks in common everyday lighting conditions as well as a 10-minute dark- and a 14-minute laboratory-based part during a ~1-hour ambulatory protocol (~300-360 samples in total) while their approximate corneal spectral irradiance and pupil size were measured. Given the 10 second sampling interval, each task and light condition took at least 60 seconds to ensure collecting enough valid samples per condition. Details about the tasks and the experimental procedure are given in Supplementary Table 1.

#### Light settings

During the natural conditions, light sources included fluorescent room lighting, LED lighting (room lighting, laptop screen) and sun light through the windows as well as outdoors. Participants moved in and between indoor and outdoor environments and engage in a range of everyday tasks while illuminance ranges from ~5 lux to many thousands of lux (upper end depending on weather conditions). Example scenarios are shown in Fig. 1E-F.

In the laboratory-based part, the light source was a vertical front panel (220 cm width, 140 cm height) mounted 80 cm above floor level. It consists of 24 LED panels (RGB + White), each containing 144 LEDs (total of 3456 LEDs; peak emission wavelengths at maximum intensity for RGB primaries: blue – 467 nm, green – 527 nm, red – 630 nm). The panel is covered with a diffuser film. Lighting scenarios were set up using DMXControl (Version 2.12.2) software (DMXControl Projects e.V., Berlin, Germany). Participants were seated on a chair in front of the light panel for 14 minutes (~75 cm distance to the eyes, height varying with participants’ height, ~90-140 cm) and instructed to gaze at a cross in the middle of the panel. The light setup comprises 4 spectrally different 3-minute phases of light (appearance: “red”, “blue”, “green”,” “white”). The 4 phases are randomized in order and embedded in 20 second dark phases (<0.2 lux before and after each light phase). Each 3-minute light phase contains 3 different illuminance levels (10 lux, 100 lux and 250–1000 lux) in ascending order, each level lasting 1 minute. The “red” and “blue” conditions reach their maximum illuminance at 480 lux and 250 lux, respectively, whereas “green” and “white” reach 1000 lux. In total the light setting adds up to 13 minutes and 40 seconds duration. The laboratory setup in shown in Fig. 1F.

#### Calibrated spectroradiometric measurements

Irradiance spectrum in the approximate corneal plane was recorded with a calibrated research-grade commercial small-footprint spectroradiometer (STS-VIS, Ocean Insight Inc., Oxford, UK; Fig. 1A), measuring in the range of 350-800 nm with a wavelength accuracy of ± 0.13nm and optical resolution of 6 nm (100 µm slit). The STS measures with a signal-to-noise ratio of >1500:1 and dynamic range of 4600:1. Integration times will be pre-set to range from 10 µs – 5 s. Attached to the STS was a direct-attach cosine corrector (CC-3-DA, diameter: 7140 µm) with 180° field of view.

The spectroradiometer was covered by a customized 3-D-printed case (Fig. 1A), adjustable in angle and mounted on a wearable, padded headband, which can be adjusted for different head sizes. The mount was centred on the forehead, such that it is approximately conjoint with the corneal plane and in the direction of gaze of the observer. The vertical angle of the spectroradiometer lens was set pointing 15° below the horizontal at default for every participant, in correspondence to typical outdoor gaze angle [50]. Irradiance spectra were measured every 10 seconds and stored on a custom-made, portable Raspberry Pi-based microcontroller.

#### Pupil size measurements

An infrared video-based eye tracker (Pupil Labs GmbH, Berlin, Germany; Fig. 1A) was used to collect still photographs of the participant’s pupil at the same time as the irradiance spectra. These still photographs (Fig. 1B) were then subjected to a 3D model estimated during a calibration procedure supplied by the eye tracker manufacturer. Images from the camera were polled using OpenCV and stored on the same microcontroller as the spectroradiometric measurements. The 3D model [51] assumes that the 3D pupil can be modelled as disk which in each moment is tangent to a rotating eye sphere. Elliptic pupil contours are extracted using a pupil segmentation and the center of the eye sphere is estimated using a nonlinear optimisation procedure (see below under “Pupil size estimation” in the “Data analysis” section). In the 3-minute-long video-based calibration procedure, the investigator verified that the 3D model could detect pupil size reliably during various head and eye movements and positions.

### Data analysis

#### Pupil size estimation

Absolute pupil diameter in millimetres was estimated from the raw images using the Pupil Labs software (release version 1.15.71, https://github.com/pupil-labs/pupil/releases). The parameters for the intrinsic 3D model are derived in the calibration stage and then applied on the pupil image sequence, yielding pupil diameter in millimetres.

#### Processing of spectral data

The spectral data reported by the spectroradiometer were given in irregular wavelength spacing and will be resampled to 1 nm intervals between 380 and 780 nm (visible range) using energy-preserving interpolation. From the spectral data we calculated the melanopic irradiance and melanopic Equivalent Daylight Illuminance^3^ (mEDI) using melanopsin spectral sensitivity curve obtained from the recent CIE S026/E:2018 standard [9] incorporating a previously defined template-derived spectral sensitivity or melanopsin [52]. In addition to the melanopic irradiance and mEDI, we also calculated the photopic illumininance in lux based on the CIE 1924 photopic luminous efficiency curve.

#### Data quality checks

We implemented the following data quality checks:

- *Lack of good-quality fit.* The pupil size estimation method will in some cases not be able to find a pupil, due to extreme angles of gaze, or partial covering by eye lashes. Any images in which a pupil cannot be found or yields low detection confidence (< *0.6*, as computed by the Pupil Labs software) was simply excluded. In a test run of the experiment, we found that ~30% of all images have this problem. This number might seem high at first, but it is important to remember that participants are performing tasks in the real world and therefore will blink and move their eyes.
- *Pupil size screening.* Since the natural pupil can only vary in a maximum range between approx. 1 and 9 mm, any estimated pupil size (due to misestimation) outside of this range was excluded.
- *Proportion of excluded data.* For each participant, the proportion of excluded data points from the entire sample will be reported. Initially, we set the following threshold: If less than 50%^4^ of data points are included, corresponding to a mere recording time of ~30 minutes, we will exclude that participant altogether.
- *Positive control and quality of pupillometry data*. We confirmed that our measurement procedure was sensible and provide upper-bound estimates for data quality by conducting laboratory-based measurements of the dark-adapted pupil size as well as twelve different conditions of light-adapted pupil size early in the protocol (for details see section “light settings” and Figure 1).

#### Recorded demographic and ancillary information

We recorded ancillary information to describe the sample. Recorded covariates include gender, handedness, BMI, vision correction, date and time of experiment, time since waking up, sleep duration, acute and habitual caffeine consumption, chronotype, acute sleepiness, iris colour and weather during the experiment. Except for the last three, these covariates were retrieved via an online questionnaire which is filled out just before and during experiment protocol.

- Iris colour and weather were rated by the experimenter. The former based on a 3-point Likert-scale item using iris colour categories “brown”, “hazel-green” and “blue” [53], rated before the begin of the protocol and the latter on a self-created 4-point Likert-scale item, incorporating the options “overcast”, “cloudy”, “partly cloudy” and “sunny”, rated at the beginning of the outdoor part of the protocol (timestamp 16 in Supplementary Table 1).
- Acute sleepiness was estimated with a German paper-pencil version of the Karolinska Sleepiness scale [54], both at the beginning and end of the protocol (timestamp 1 and 29 in Supplementary Table 1). The 9-point Likert-scale item ranges is labelled on all odd steps.
- Both the habitual and acute caffeine consumption questionnaires were developed based on median values of the caffeine content ranges in mg for typical caffeinated beverages and food items [55]^5^. Each questionnaire comprises 18 items (15 on beverages, 2 on chocolate and 1 for caffeine pills). Each item asks the participant to indicate how many portions of the caffeinated item were consumed on a scale from 0 (no) portion to 8 portions. Every item portion matches a specified typical portion size in ml or g and a caffeine content in mg. While the “habitual” questionnaire is asking for intake on an average day, the “acute” questionnaire queries the same items with regards to the last 6 hours. For both habitual and acute intake, an absolute sum and a relative sum value of caffeine in mg (divided by weight in kg) are computed for every participant. The two questionnaires were filled out during the protocol as part of a “laptop task” (timestamp 12 in Supplementary Table 1).
- Chronotype was queried with a German translation of the µMCTQ, an ultra-short version of the Munich ChronoType Questionnaire [56]. It comprises six core questions, allowing for a quick and efficient assessment of chronotype. The indicating value for chronotype is computed as the midpoint of sleep on work-free days corrected by potential sleep debt (MSF_SC_). Sleep duration and time since waking up are assessed from answers to four simple open-ended questions. Participants indicate the times of falling asleep and waking up concerning the night before the experiment as well as the duration of potential naps took on the day of and before the experiment. Sleep duration is computed by subtracting time of waking up from time of falling asleep and adding nap duration. Time since waking up is computed as the time starting to fill out the questionnaire minus the time of waking up. Both the µMCTQ and the questions concerned with sleep duration and time since waking up were filled out early in the protocol as part of a “laptop task” (timestamp 4 in Supplementary Table 1).
- Gender, handedness, BMI (computed from weight and height) and vision correction (“no correction”, “minus correction” or “plus correction”) were assessed via simple demographic items as part of the online questionnaire, filled out just before the beginning of the protocol. Date and time of the experiment were logged by the start of filling out that online questionnaire.

These parameters were recorded to describe the sample and not used in our confirmatory hypotheses tests^6^.

#### Analytic strategy

We used a linear mixed model framework to examine our three confirmatory hypotheses and examine evidence for each of our three hypotheses using Bayes Factor [57]. The general logic in this approach is that a ‘full’ model containing a factor under consideration is compared to a ‘null’ model missing that factor using the likelihood ratio between the two models. If the factor affects the variable in question (pupil size), this will be reflected in higher Bayes Factors. We consider a BF of 10 as “strong” evidence, following standard categorisations [58]: 1 < BF < 3 – anecdotal evidence, 3 < BF < 10 – moderate, 10 < BF < 30 – strong evidence, and BF > 30 – very strong evidence, BF > 100 – decisive evidence^7^. We assess three confirmatory hypotheses independently and convert the Bayes Factors obtained to this scale of evidence. The analysed relationship between pupil size and light refers to simultaneously taken samples of those variables.

Our first confirmatory hypothesis (CH1) tests whether melanopsin sensitivity-weighted irradiance predicts our pupil size. The formulation of the full model in Wilkinson-Rogers notation [59] is as follows:

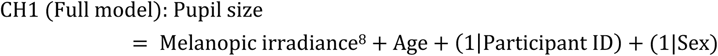

This full model is compared to the null model not containing melanopsin irradiance:

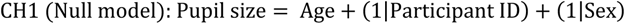

The second confirmatory hypothesis tests if melanopsin sensitivity-weighted retinal illumination better predicts pupil size than the weighted sum of L- and M-cone-weighted retinal illumination (CIE 1931 *Y*, photopic illuminance). We express this full model as:

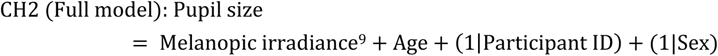

Here, the null model is:

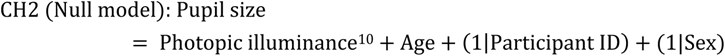

Our third confirmatory hypothesis will test if pupil size changes as a function of age, with higher age associated with a smaller pupil size. We express the full model as:

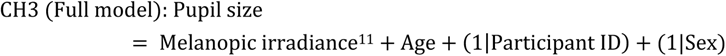

Here, the null model is:

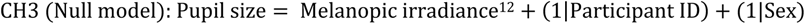

#### Power analysis

There were no prior data allowing us to conduct a meaningful power analysis. The largest (to our knowledge) study examining the relationship between pupil size and illuminance [28] only used one spectral stimulus (scaled to different illuminances). We considered the question of power as follows. In hypothesis CH1, we aimed to confirm the light-level dependence of pupil size. Unless a participant has an undetected neurological or retinal condition leading to immobility or paradoxical behaviour of the pupil, the light-level effect will be present in all individuals. Therefore, the effect is nearly guaranteed to exist in a given individual. In hypothesis CH2, we examined whether melanopic irradiance was a better predictor for pupil size than photopic illuminance. From our previous work, we know that the spectral sensitivity of pupil size is best described using melanopsin rather than photopic luminosity [15]. In the laboratory part of the experiment, we near-recreate these conditions using narrowband LED lights. Regarding hypothesis CH3, we were confident in the study’s power to detect effects of age on pupil size in an age-diverse sample (*N=96)*. Very stable effects of age on pupil size were shown in a comparable sample (*N=91,* [28]). Our sample size (96 individuals) was governed by our resource limitations.

#### Data collection termination rules

Data will be collected until our resource limit (96 participants and study personnel hours) is reached.

### Data accessibility

All code and data are publicly available as part of this publication. Code, raw and processed data (except code to drive the acquisition device, which is proprietary) are available on GitHub (https://github.com/tscnlab/LazarEtAl_RSocOpenSci_2023) under the <u>MIT</u> license. Raw data (https://doi.org/10.6084/m9.figshare.24230848.v1) and supporting material (https://doi.org/10.6084/m9.figshare.24230890.v1) are available on FigShare under the <u>CC-BY 4.0</u> license. The in-principle accepted Stage 1 protocol can be found here: (https://osf.io/zrksf/).

### Ethical approval

Ethical approval has been granted from the cantonal ethics commission (Ethikkommission Nordwest-und Zentralschweiz, project ID 2019-01832, see Supporting materials on <u>FigShare</u>). All current mandatory measures to prevent the spread of COVID-19 in Switzerland were implemented during data collection.

### Known limitations of this registered report

We acknowledge that our study is limited with respect to following aspects:

- *Other, small-scale determinants of pupil size are ignored.* In this work, we are agnostic to other determinants of pupil size, such as fatigue [60], attentional processes [61, 62], recognition [63], high-level image content [64], target detection [65], mental effort [63, 66], arousal [67] and substance use [68]. These top-down effects are expected to be transient and relatively small in effect compared to the light and age effect [2, 35]. In addition, due to the near triad comprising convergence eye movements and ocular accommodation, viewing distance affects pupil size due to a shared neural pathway [1].
- *Estimation of retinal illuminance in natural vision.* In this study, we were using an irradiance sensor near the corneal plane, placed on the forehead and adjusted to typical outdoor gaze angle (15° below the horizontal, [50]). Given the geometry of the sensor and human head, this is the closest we can get to measuring corneal illuminance. It is impossible to estimate retinal illuminance using sensors in the real-world under freely changing geometries and gaze positions.
- *Light history.* In the present study, we are omitting effects of prior light exposure (e.g. before the experiment) and light sequence effects during the protocol on pupil size. The protocol procedure of the field experiment is held constant across participants (except for the randomized light phases in the laboratory-based part of the experiment)
- *Omitting scene radiance.* In the present study we did not measure spectral radiance but spectral irradiance. By doing this, we neglect effects of variation in light level across the field of view in different light scenes. Given the limitations of the real-world setting, the technical properties and calibration of the available instruments, collecting real-world radiance data and artificially limiting the angular acceptance of our spectroradiometer is not feasible in the present study.

## Results

We carried out the experiment according to the Stage 1 protocol, which was accepted in principle (https://osf.io/zrksf/). All LMM analyses were performed in R (v. 4.3.1) [69] using the BayesFactor package (v. 0.9.12-4.5) [70]. The “ggplot2” package (v. 3.4.3) [71] was utilized for data visualization. More details are available in the supplementary information.

In the following, we first report participant characteristics and quality check analyses. We then present the results of our confirmatory hypotheses using a final sample of 83 participants and log_10_-transformed light data. For transparency and completeness, we also report the results of our hypothesis tests in the linear, non-transformed light data despite the violation of linear assumptions both in the full (n=83, 75% data loss threshold) and in the reduced sample of 63 participants (50% data loss threshold). Finally, we summarise our post-hoc analyses.

### Participant recruitment

Details of the recruitment process are presented in the study flow diagram (Suppl. Figure 1). Following our recruitment plan of the approved Stage 1 registration, data was collected until we reached our resource limit. Due to difficulties filling the older age groups during the COVID-19 pandemic, we later opened recruitment to more than 16 per age group to achieve a large enough total sample. In total, n=113 volunteers were invited to the health and eligibility screening. Seven did not meet the inclusion criteria – five because of the visual tests and two for health reasons and medication with potential influence on pupil regulation. The remaining 106 subjects were allocated to the protocol. While n=87 completed the trials without problems with the equipment, data from n=19 volunteers’ trials were invalid due to technical problems (cable issue, camera slippage or software/format errors) and had to be excluded. A further four participants were then removed before the final analysis because more than 75%^13^ of their pupil data were of insufficient quality (see *Data quality checks*), leaving n=83 for the final analysis. Given the resource limitations mentioned above, this sample remained as the final sample for statistical analysis.

### Participant characteristics

Table 1 presents sample characteristics based on self-report separately for participants who were included (n=83), excluded after the study (n=23), and excluded before the study (n=7). Among the 83 volunteers included in the final analyses (43 female, 40 male; *M*_age_ = 35.70, *SD*_age_ = 17.16; *M*_BMI_ = 22.96, *SD*_BMI_ = 3.47), age groups were not balanced as planned initially (16 per group), and the final sample is skewed towards younger age, with 30 included volunteers aged 18-24 years and five included volunteers over 64. This is also depicted in Figure 2, where the included sample is stratified by sex. Notably, the volunteers excluded before the trial were clearly older on average and with higher BMI (*M*_age_ = 52.00, *SD*_age_ = 21.54, *M*_BMI_ = 26.39, *SD*_BMI_ = 5.79).

A total of 40 included participants took part in the experiment during summer, 20 during winter, 16 during autumn and 7 during spring. Weather conditions varied between light rain (11%), very cloudy (19%), cloudy (17%), somewhat cloudy (13%) and sunny (40%). Differences in weather conditions during the trial were associated with different light intensity distributions. Figure 3 depicts the density of light intensity (mEDI) across the whole trial (upper panel) and stratified across the weather conditions (lower panel). Notably, in sunny weather, there is a higher probability density of mEDI values over 1000 lx than in all other weather conditions. Additional participant information like chronotype, sleep duration, start time of the experiment and acute and habitual caffeine consumption are given in Suppl. Table 2.

**Figure 3:**
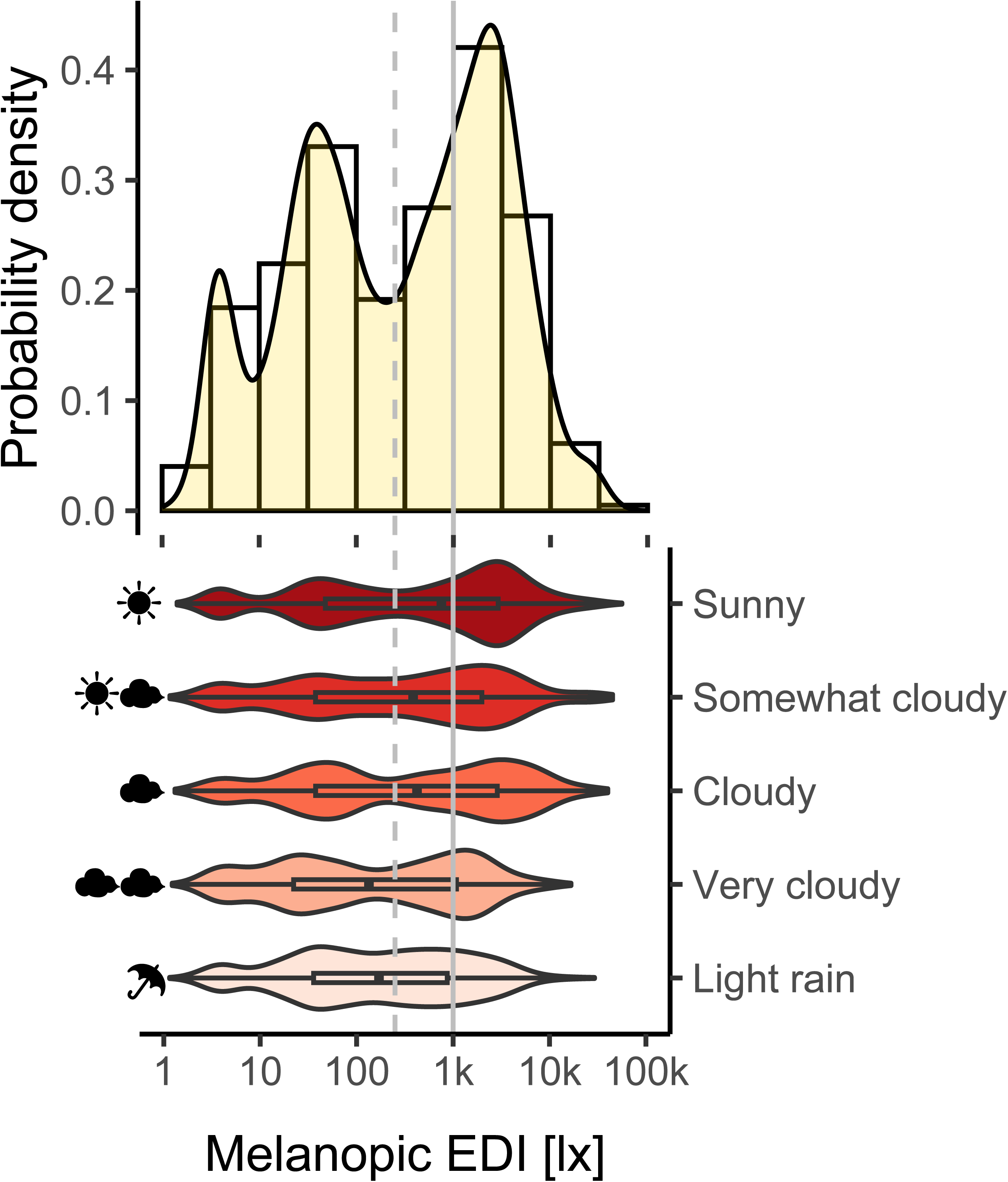
Density of light intensity (melanopic EDI) across all observations in the field conditions (n=83, top panel) and stratified across the weather conditions (bottom panel). The dotted and solid vertical grey lines indicate intensities of 250 lx mEDI and 1000 lx mEDI, respectively. Bin breaks in the histogram (top panel) correspond to half log10-unit steps of mEDI. Box plots in the density violin plots (bottom panel) include the median (black square) along with the interquartile range (IQR, black rectangle) under each weather condition. The “whiskers” mark the values below the 1^st^ and beyond the 4^th^ quartile. Sunny weather is associated with a larger proportion of >1000 lx mEDI values.

### Data quality checks

The quality checks specified earlier in this report were applied and further supplemented by a check of linear assumptions for the hypothesized mixed linear models.

#### Proportion of excluded data

Any pupil images yielding low detection confidence (<0.6) or exceeding the maximum range (<1 or >9 mm) were replaced by missing values, as were saturated spectroradiometer samples (after quick changes from dark to bright). Consequentially, we calculated the proportion of excluded data points for each participant (see Suppl. Table 3).

Initially, we had set a threshold of maximally 50% data loss per subject. Otherwise, we planned to exclude that subject altogether. As our dataset revealed a higher proportion of pupil data exclusion than expected from the pilot runs, this would have amounted to excluding an additional n=24 (out of n=87) participants from the final analysis (cf. Figure 4). Thus, to address high participant attrition, we decided to additionally determine a second adjusted threshold of “acceptable” data loss proportion per participant after Stage 1 in principle acceptance (IPA).

**Figure 4:**
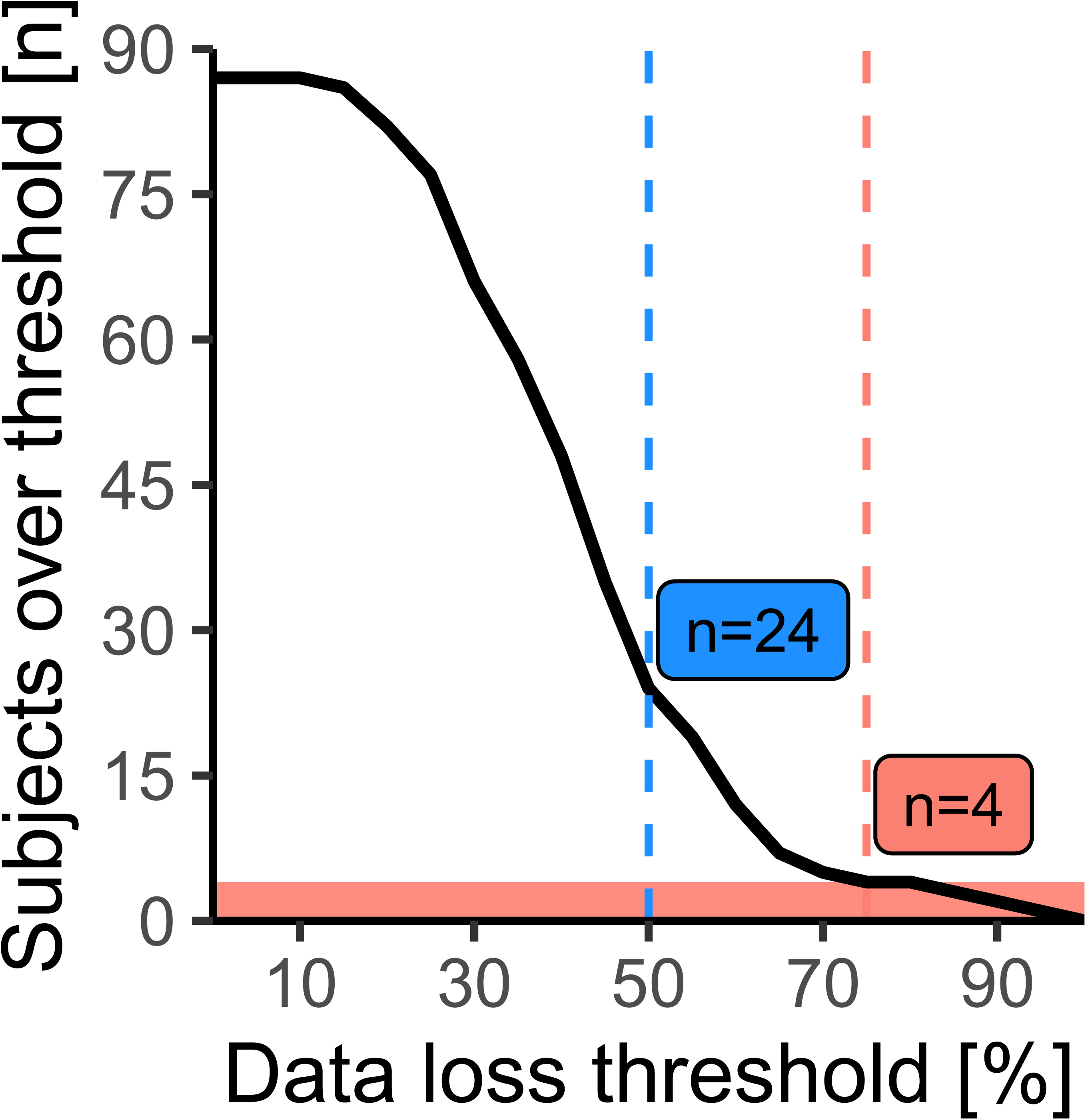
Excluded participants as a function of possible data loss thresholds (black line). The initial data loss threshold (50%) is marked by the dashed blue vertical line, yielding n=24 excluded participants. The adjusted threshold (dashed magenta line) was placed at the location over 50%, which marks the smallest increase of excluded participants (+0, from 75 to 80%)) across possible thresholds in 5% steps, yielding n=4 excluded participants and retaining n=83.

We determined a second sensible “data loss threshold” in a data-driven analysis of a “scree-like” plot. The criterion was to find the location beyond the 50% threshold, where increasing the threshold in 5% steps would lead to a minimally increased number of excluded participants. As depicted in Figure 4, this procedure resulted in adopting a second data loss threshold of 75% excluded data points per participant, with n=4 (out of n=87) exceeding the new threshold with values of 81.60%, 85.48%, 91.67 %, and 95.65%.

In the remaining dataset of n=83 (all conditions), a total of 17199 observations with good-quality pupil and light data were retained. The full dataset is separated into “field data” (10082 valid observations), “dark data” (2871 valid pupil observations), and “lab data” (3698 valid observations). There were also residual “transition samples” (548 valid observations) taken between the laboratory and field conditions that were not used in the analyses because they could not be clearly assigned to either condition.

Data loss per included participant ranges from 10.82% to 70.64% as listed in Suppl. Table 3. Furthermore Suppl. Tables 3 and 4 display a summary of minimum, median and maximum values for mEDI and pupil size, separated for field and positive control data (laboratory and dark adaptation conditions), respectively. Only observations from the field condition with retained pupil and light data were used for confirmatory hypothesis testing.

#### Linear Regression Assumptions

We discovered a peculiarity in the originally formulated linear mixed model analyses after receiving Stage 1 IPA and collecting data. When predicting pupil size with linear light intensity data (mEDI/melanopic irradiance and photopic illuminance), these variables violate the assumptions of linear regression models, specifically the assumptions for linearity, homogeneity of variance and collinearity (see Suppl. Figure 2 B, C and E). When light data were log_10_-transformed prior to inclusion in the linear models, the requirements were approximately met (see Suppl. Figure 3), suggesting transformation. We additionally tested whether the transformation improved the fit in the LMM, comparing the model with log_10_ transformation against a null model with linear light data in the same procedure as the hypothesis tests in the pre-registered analysis. As shown in Suppl. Table 5, the resulting likelihood ratio clearly favours the model with log_10_ transformed light data (BF_10_=5.678918e+996 ±3.55% pe), corresponding to “decisive evidence” in favour of a log_10_-transformation. For completeness and transparency, we report the results of our hypothesis tests both with transformed and non-transformed data.

#### Positive control data

Two conditions were included in the experiment as a manipulation control: First, a 10-minute dark adaptation to test whether the age effect is generally present. Second, a laboratory light condition in which different light colours allow stronger dissociation between photopic illuminance and mEDI values (see *Light settings*), to confirm that mEDI determines pupil size in the light-adapted pupil and is superior as a predictor compared to photopic illuminance.

The recorded light intensities from the laboratory condition, contrasted with the field conditions, are shown in Figure 5. In comparison to the extremely high positive correlation between photopic illuminance and mEDI under uncontrolled field conditions (r=.999, see panel B), the laboratory condition yielded a clearly lower positive correlation (r=.309, see panel A). We tested confirmatory hypotheses CH1 and CH2 with the linear models specified in stage 1 of this report with data from these low-correlation laboratory conditions. Both hypotheses are supported by decisive evidence (BF_10_>100). This is the case for the full sample (n=83, 70% threshold) as well as the reduced sample (n=63, 50% threshold) and with log_10_-transformed as well as linear light data (see rows 1-4 of Suppl. Tables 6, 7, respectively).

**Figure 5:**
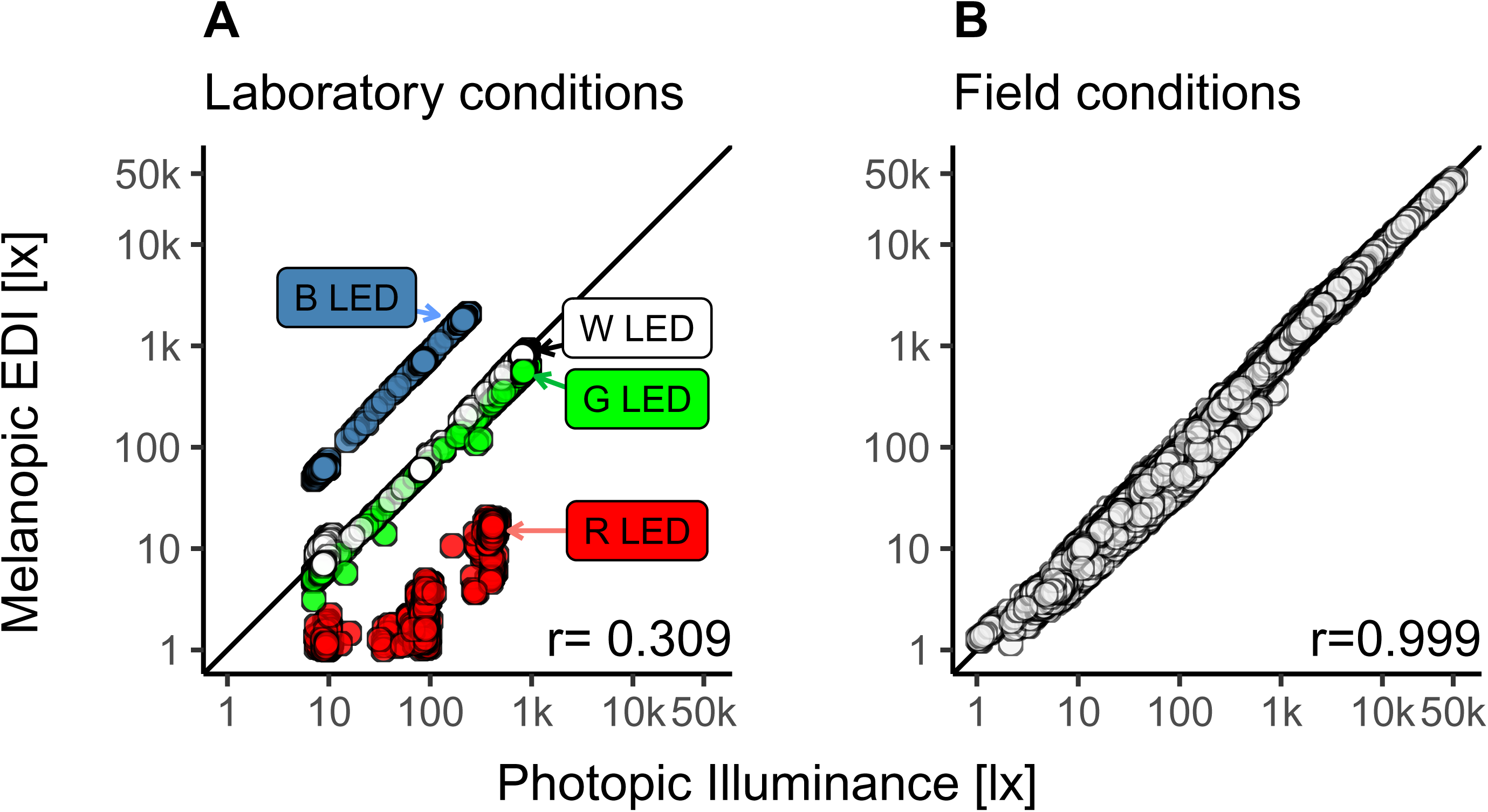
(***A***-***B***) Relationship between melanopic and photopic light units contrasted between laboratory conditions (***A***) and field conditions (***B***). Pearson correlation coefficients quantify the association. The laboratory data clearly show a dissociation between the light intensity units, whereas, under field conditions, these units are highly correlated.

Testing the age effect under dark conditions, we compared a model including age as a predictor vs. the null model without it. The third row of Suppl. Tables 6 and 7, present the tested models and results in the full sample (n=83, 70% threshold) and reduced sample (n=63, 50% threshold), respectively. Both provide decisive evidence (BF_10_>100) for age as an influencing factor on pupil size during dark adaptation.

### Confirmatory analyses

#### Pupillary light reflex

Hypothesis CH1 proposed that pupil size is dependent on the (melanopsin sensitivity-weighted) light level under naturalistic steady-state conditions. Our results from the LMM analysis (n=83, 70% threshold) provide decisive evidence (BF_10_>100) for that hypothesis, both when using the log_10_-transformed mEDI data (cf. Table 2, row 1) and the linear mEDI data in the full model (cf. Suppl Table 8, row 1). Notably, the likelihood ratio turns out to be magnitudes higher after log_10_-transformation compared to without it (BF_10_=1.543255e+1288 ±2.20% pe vs. BF_10_=2.642936e+291 ±2.78% pe, respectively).

Considering the reduced dataset (n=63, 50% threshold), the hypotheses tests produce qualitatively similar results, yielding decisive evidence for the full model and CH1 (BF_10_>100), again for both the log_10_-transformed and linear data (cf. Suppl. Table 9, row 1 and Suppl. Table 10, row 1). Likelihood ratios are larger in the full sample (see above) compared to in the reduced sample (BF_10_=2.404706e+1076 ±2.24% pe and BF_10_=1.031709e+242±2.24% pe, respectively).

#### Predictive superiority of melanopsin sensitivity weighted measures

In hypothesis CH2, we tested whether mEDI was a better predictor of steady-state pupil size than photopic illuminance. The LMM results (n=83, 70% threshold) support this hypothesis with decisive evidence (BF_10_>100), both using the log10-transformed light data (see Table 2, row 2) and the linear mEDI data (see Suppl. Table 8, row 2). Once more, the likelihood ratio turns out to be several orders of magnitude higher after log_10_ transformation than without (BF_10_=1.458261e+21 ±2.13% pe vs. BF_10_=10708989894 ±3.91% pe, respectively).

Looking at the reduced dataset (n=63, 50% threshold), the hypothesis tests yield qualitatively equivalent results, providing decisive evidence for the full model of CH2 (BF_10_> 100), again for both the log_10_-transformed and linear data (see Suppl. Table 9, row 2 and Suppl. Table 10, row 2). The Bayes factors are again larger in the full sample (see above) than in the reduced sample (BF_10_=1.782022e+19 ±2.06% pe and BF_10_=1921990 ±1.93% pe, respectively).

#### Age effect in the light-adapted pupil

Hypothesis CH3 stated that light-adapted pupil size decreases (on average) with increasing age, a phenomenon also termed “senile miosis”. Results from the LMM (n=83, 70% threshold) support the notion that age is a significant factor for pupil size with decisive evidence (BF_10_>100). This is the case when including log_10_-transformed mEDI data (see Table 2, row 3) or linear mEDI data in the models (see Suppl. Table 8, row 3). The likelihood ratio comparing the full and null models, including the log_10_-transformed data, was ~20 times higher than the BF of the model comparison without transformation (BF_10_=46076.190 ±2.17% pe vs. BF_10_=2221.5 ±2.66% pe, respectively).

In the reduced data set (n=63, 50% threshold), the hypothesis test with the full model of CH3, including age and the log_10_-transformed mEDI data, leads to the same interpretation: Decisive evidence in favour of CH3 (BF_10_=124.465 ±2.12% pe; see Suppl. Table 9, row 3). However, the likelihood ratio for the full model of CH3, including age and the linear mEDI data, falls below 100 (BF_10_=13.992 ±1.89% pe, see Suppl. Table 10, row 3) and is therefore interpreted as “strong evidence” (BF_10_>10) for CH3 instead.

### Post-hoc Analyses

As an extension of our confirmatory hypotheses, we visualized the direction and magnitude of the tested age effects on pupil size, inspired by the work of Winn et al. [28]. We further performed explorative analyses testing sex, iris colour and reported acute and habitual caffeine consumption (relative to body weight) as potential influence factors on light-adapted pupil size under real-world conditions. These analyses were not included in our Stage 1 submission.

#### Visualising the age effect

Providing typical case data, Figure 6 depicts pupil size data from a young (18 years of age) and an old participant (87 years of age), plotted against mEDI, including dark adaptation and field data from the experiment. As can be seen from the maximum to minimum range (given by the coloured horizontal lines) and from the regression equation, the older participant has a clearly smaller baseline pupil diameter (β_0_=3.82 mm) and magnitude of pupillary light reflex (β_1_=−0.462 mm per log-unit mEDI) compared to the young subject (vs. β_0_=6.26 mm and β_1_=−0.997 mm per log-unit mEDI). Interestingly, the variation in pupil size within similar light intensities decreases markedly towards the high mEDI intensities.

**Figure 6:**
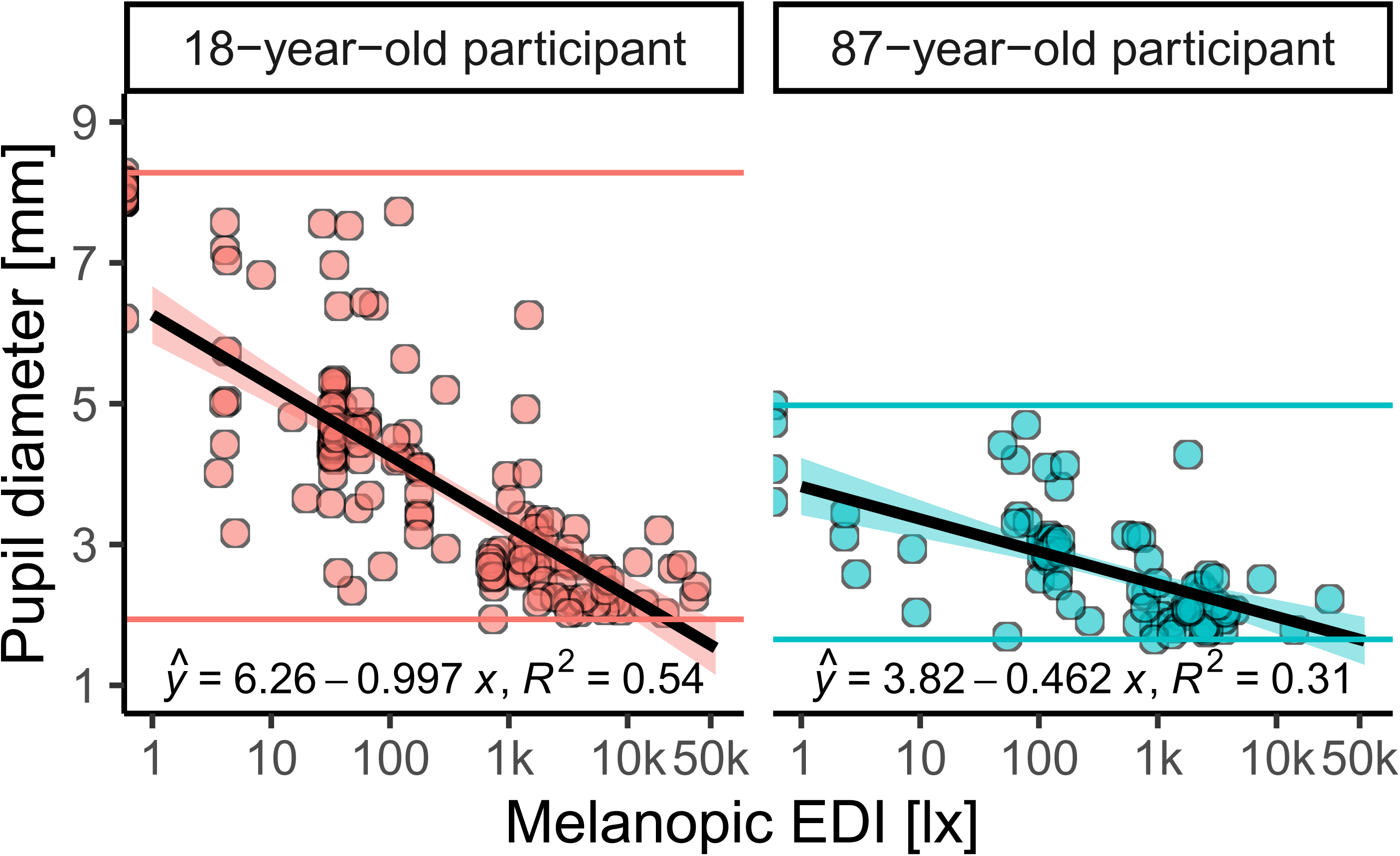
Age comparison of typical pupil size data as a function of melanopic Equivalent Daylight Illuminance (mEDI). The scatterplots show data from the field and dark conditions of a young (18 years of age, left panel) and old participant (87 years of age, right panel). The regression line and equation demonstrate the linear relationship between log_10_-transformed mEDI values in lux and pupil diameter in mm. The coloured horizontal lines mark each subject’s minimum and maximum pupil diameter values. A reduced pupil dilation in the lower light intensities is visible for the older subject.

Inspired by Winn et al. (cf. Figure 2 in [28]), Figure 7 visualises the direction and magnitude of the age effect across the full sample (n=83) under different light condition clusters given in mEDI. Coloured squares depict each subject’s median pupil size values under the respective condition plotted against age in years along with the interquartile range (see grey opaque bars). The linear regression lines are fitted to the median values, while the 95% confidence limits are indicated by the grey shade around the line.

**Figure 7:**
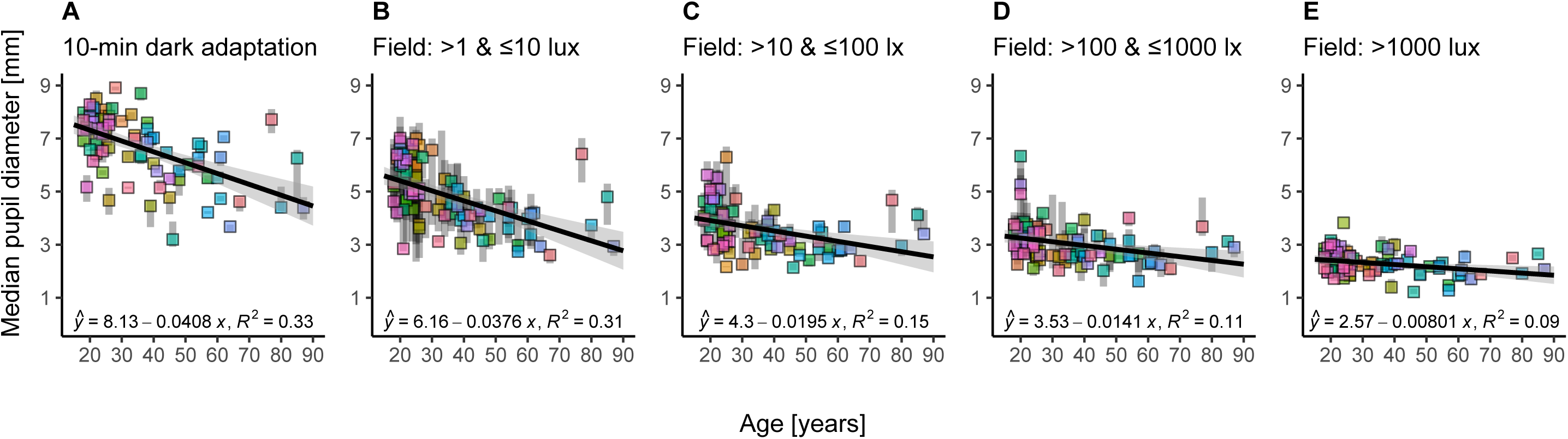
(A-E) Median values of pupil size per subject (n=83) as a function of age during different light conditions. *A* In the 10-minute laboratory-based dark adaptation condition. *B* In Field conditions between 1 and 10 lx mEDI. *C* In Field conditions between 10 and 100 lx mEDI. *D* In Field conditions between 100 and 1000 lx mEDI. E In Field conditions over 1000 lx mEDI. The coloured squares display the median values per subject along with the interquartile range (IQR, opaque grey bars). The equations and regression lines, including the 95% confidence intervals (grey line shade), demonstrate the linear relationship between age in years and median pupil diameter in mm across the different clusters of light intensities during the experiment. Field data (*B-E*) were included in the confirmatory hypothesis analysis, while dark data (*A*) was only used in the positive control analysis. The effect of ageing is reduced as the intensity of the light increases.

Figure 7 Panel A shows the age effect during the 10-minute in-lab dark adaptation (data not used in the confirmatory analyses), while panels B-E depict the age effect under field conditions in log_10_-unit steps. In all light conditions, we see an apparent reduction of pupil size with increasing age (on average), consistent with the beta values in the regression equations given in each panel. Interestingly, the slopes and intercepts of the age effect decrease with increasing light intensity (mEDI) across all intensity clusters, as shown in Figure 8 A & B, along with the 95% confidence intervals given as error bars.

**Figure 8:**
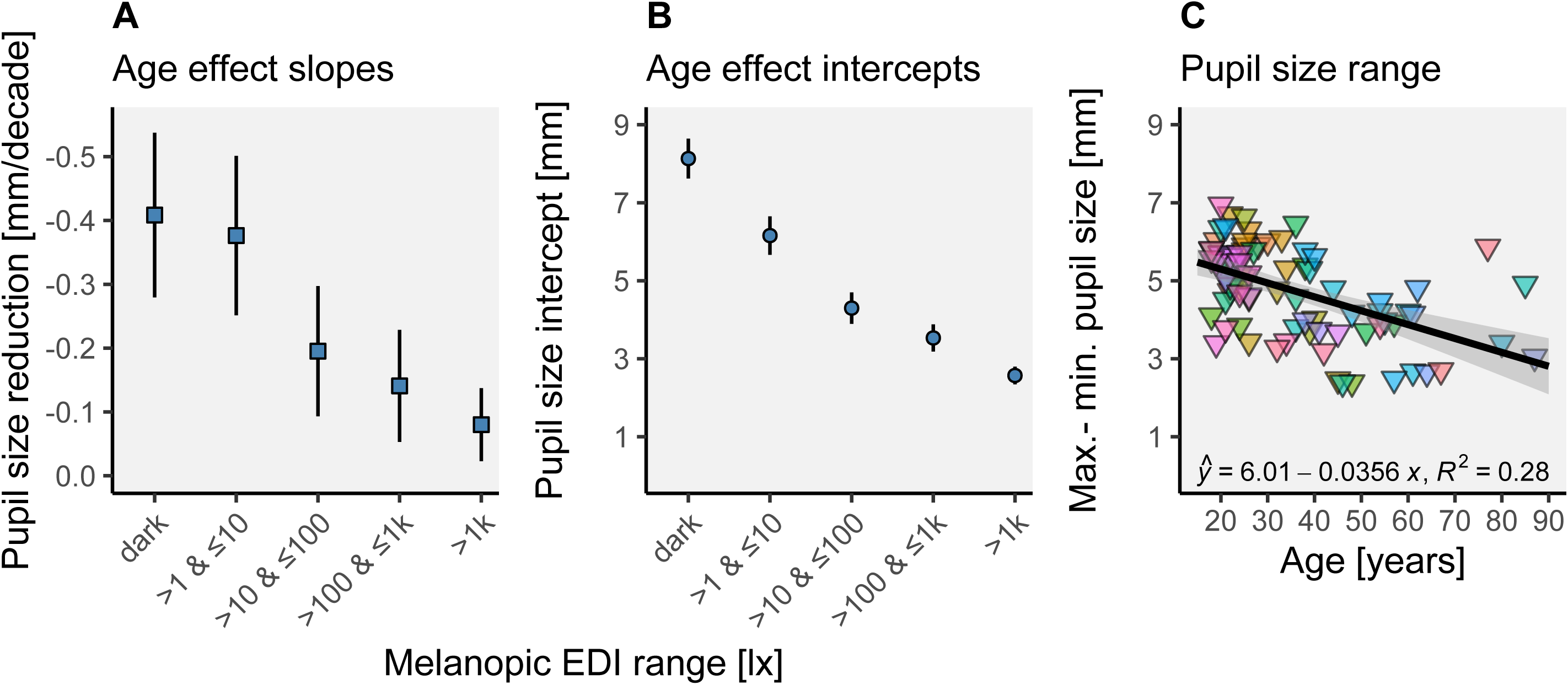
(A-B) Slopes (*A*) and Intercepts (*B*) of the linear regressions for median pupil size as a function of age (in decades), plotted per light intensity cluster. Error bars give the 95% confidence intervals of the parameters. The slopes and intercepts of the age effect and their 95% confidence intervals continuously decrease with increasing light intensity (cf. Figure 7). *C* Pupil size range (maximum-minimum diameter in mm) as a function of age in years. The equation and regression line, including the 95% confidence interval (grey line shade), demonstrate the linear relationship between age in years and pupil size range in mm.

The magnitudes of the age effect (summarized in Figure 8 A) are highest in the dark-adapted pupil with pupil size reduction of β_1_=−0.408 mm per decade (CI_95%_=[−0.538, −0.279]). Under field conditions between 1 and 10 lx mEDI, age-driven pupil size reduction amounts to β_1_=−0.376 mm per decade (CI_95%_=[−0.501, −0.251]), between 10 and 100 lx mEDI, it amounts to β_1_=−0.195 mm per decade (CI_95%_=[−0.297, −0.093]), and between 100 and 1000 lx mEDI the age effect results in a reduction of β_1_=−0.141 mm per decade (CI_95%_=[−0.229, −0.053]). Finally, in the brightest light conditions (>1000 lx mEDI), the age effect estimation is smallest with β_1_=−0.080 mm per decade (CI_95%_=[−0.137, −0.023]).

The intercepts of the age effect (Figure 8 B) range from fully dilated pupils at β_0_=8.13 mm in darkness (CI_95%_=[7.62, 8.64]) to fully constricted pupils at β_0_=2.57 mm (CI_95%_=[2.35, 2.80]) under bright light field conditions (>1000 lx mEDI). For the intermediate steps, field conditions between 1 and 10 lx mEDI yield a pupil size intercept of β_0_=6.16 mm (CI_95%_=[5.66, 6.65]), field conditions between 10 and 100 lx mEDI, yield β_0_=4.30 mm (CI_95%_=[3.90, 4.70]), while field conditions between 100 and 1000 lx mEDI results in β_0_=3.53 (CI_95%_=[3.19, 3.88]).

However, we find considerable inter-individual variability in pupil size, even among participants of similar age, at all mEDI levels. With increasing light intensity, this variability is reduced, showing a close resemblance to the laboratory data of Winn and colleagues [28]. It is also worth noting that the individuals’ interquartile ranges of pupil size are lower during controlled ten-minute in-lab dark adaption but, under field conditions, decrease as the light intensity increases (cf. IQR in Figure 7A, B and E)

Figure 8 C depicts the maximum to minimum pupil size range in mm plotted against age in years across all subjects. Equivalent to pupil size under different light intensities (see Figure 7 A-E), there is an apparent negative linear effect on pupil size range with β_1_=−0.356 mm per decade (CI_95%_=[−0.482, −0.230]) and an intercept of β_0_=6.01 mm (CI_95%_=[5.51, 6.51]).

#### Exploratory hypotheses

According to the same LMM procedure as in the confirmatory analyses (see *Analytic Strategy* under *Materials and Methods*), we tested sex, iris colour, and caffeine consumption (habitual and acute) as potential factors influencing light-adapted pupil size under natural conditions in the full sample (n=83, 70% threshold). We derived the following exploratory hypotheses:

- EH1: Sex differences in light-adapted pupil size are present under real-world conditions.
- EH2: Light-adapted pupil size varies as a function of iris colour under real-world conditions.
- EH3: Light-adapted pupil size varies as a function of habitual caffeine consumption (relative to body weight) under real-world conditions.
- EH4: Light-adapted pupil size varies as a function of acute caffeine consumption (relative to body weight) under real-world conditions.

As summarised in Table 3, along with the corresponding model comparisons in Wilkinson-Rogers’ notation [59], none of the additional factors tested for influence on light-adapted pupil size proved to be supported by evidence from our sample. We found strong evidence in support of the null models (BF_10_ <0.1) for sex and iris colour, suggesting that there are no sex differences in light-adapted pupil size under real-world conditions (BF_10_=0.070 ±1.44% pe, see Table 3, row one) and that pupil size does not vary as a function of iris colour (BF_10_=0.017 ±2.86% pe, see Table 3, row two). We also observed moderate evidence in support of the null models (BF_10_ <1/3) for habitual and acute caffeine consumption per body weight [mg/kg], suggesting that neither habitual (BF_10_=0.131 ±2.10% pe, see Table 3, row three) nor acute caffeine consumption within the last six hours (BF_10_=0.176 ±2.10% pe, see Table 3, row four) had a significant effect on pupil size in our sample under real-world conditions.

## Discussion

Here, we conducted a real-world experimental study using a novel ambulatory setup to perform confirmatory analyses of age effects and effects of spectrally weighted illumination on light-adapted pupil size. To this end, we measured light-adapted pupil size and spectral irradiance during ecologically relevant tasks in everyday indoor and outdoor environments under naturalistic lighting conditions in a healthy, age-diverse and gender-balanced sample.

### Positive Control data

Our laboratory data analyses as positive control indicated that our measurements were valid and that our confirmatory hypotheses were supported by decisive evidence, consistent with prior in-lab work [11–14, 19–21].

However, despite our prior test runs, we underestimated data loss due to the many possible interference factors when performing ecologically relevant tasks in dynamic real-world conditions. Therefore, we adjusted our initial data loss threshold to avoid high participant attrition. In our confirmatory analyses, we consistently reported both the full and a reduced dataset, showing that while the interpretation did not change, the likelihood ratios in support of the hypotheses were higher in the larger sample.

### Log_10_-transformation of light intensity

To deal with the unanticipated violation of linear regression assumptions when including the light data as predictors, we log_10_-transformed them, which resolved the issue. However, this step was not planned in our Stage 1 Registered Report, which led us to always report both transformed and untransformed data in our confirmatory analyses for full transparency.

Besides statistical reasons, predicting pupil size employing a log_10_-transformed light unit also conceptually makes sense. Light exposure under naturalistic conditions spans several orders of magnitude (up to a factor of 10^10^) [3, 72]. It has been suggested that the visual system performs some type of logarithmic compression, not least as it enables scaling of signals according to their proportions rather than absolute scale owing to the difference in solar irradiance [73, 74]. The melanopsin-encoded component of ipRGCs appears to operate in a log-linear fashion, there by encoding irradiance across several orders of magnitude [75, 76]. This is consistent with the finding in our dataset that the log_10_ transformation clearly improves the pupil size prediction when compared to the linear scaled data.

### Pupillary light reflex in the field

Decisive evidence from our field data confirms the presence of the pupillary light reflex under natural conditions (CH1). We expected this effect to be present in all participants unless they had an unrecognised medical condition.

We found distinct dose-response curves for the case data presented in Figure 6, with smaller pupil sizes observed at brighter conditions. The resulting curves closely resemble the model predictions based on laboratory luminance data shown in Figure 1 C [35]. Interestingly, the variation in pupil size within similar light intensities decreases markedly towards the high mEDI intensities. This finding may reflect that the pupillary light reflex dominates pupil regulation at high light intensities, whereas, at lower intensities, there is room for variation due to the other effects on pupil size, such as accommodation or emotional and cognitive responses.

### Predictive superiority of melanopsin sensitivity weighted measures

Despite the extremely high correlation between photopic illuminance and mEDI under our field conditions, the analysis revealed the clear superiority of melanopsin-sensitivity weighted measures over photopic illuminance in predicting light-adapted pupil size under our natural light and task conditions (CH2). This effect was robust across the reduced and full dataset and log_10_- transformed and linear light data. Our finding confirms evidence from laboratory studies suggesting that mean steady-state pupil size under photopic conditions is primarily controlled by the melanopsin-encoded signal [11–16].

These findings provide a clear call to action: The application of melanopsin-weighted measures (e.g., mEDI) should be extended beyond the laboratory, which is likely to lead to significant improvements in pupil size prediction compared to photopic illuminance.

### Age effect on pupil size

In line with previous laboratory findings [19–28], our data confirmed that steady-state pupil size decreases with age, even under uncontrolled real-world conditions. In addition, our exploratory results replicate that the effect is stronger in dim compared to bright light conditions [13, 24, 28, 29, 77]. Perhaps the most compelling findings are the consistent linear regression parameters between our data and the laboratory data presented in Figure 2 by Winn et al. in 1994 [28]. Both studies indicate that under dim light conditions, ageing reduces pupil size by about 0.4 mm per decade, starting at ~8 mm intercept. Surprisingly, the clustered field data also closely match the effect magnitudes in the laboratory, as reported by Winn and colleagues. In line with this pattern, our data extend to even brighter daylight conditions, where the age effect is even more reduced, and pupils are likely fully constricted. Lastly, just like in the findings of Winn and colleagues, our data also show “a substantial amount of interindividual variation in pupil size for subjects of similar age” [cf. 28p. 1134] and decreasing variation with increasing brightness.

In summary, age reduces pupil size in the real world, especially at lower light intensities, albeit with large inter-individual variability. This provides further support for considering age-related differences in lighting solutions for visual and non-visual optimisation, as smaller pupil sizes result in reduced retinal illumination. A study by Gimenez et al. [78]suggests that light-induced melatonin reduction in healthy young people might adapt to long-term reductions in short-wavelength light through filters. Whether this is the case in an older population experiencing symptoms of the ageing eye, such as lens yellowing, photoreceptor loss and pupil size reduction, remains an important research question. Prior evidence suggests that there are some compensatory mechanisms at play in ageing [79].

### Other potential factors affecting pupil size

Derived from the literature, sex, iris colour, and caffeine consumption were investigated as potential influencing factors. Previously, Abokyi et al [68] found that acute caffeine intake of 250 mg significantly increased pupil diameter from 30 to 90 minutes after ingestion. In contrast, our data show no evidence that caffeine intake “within the last six hours” or habitual caffeine consumption affect light-adapted pupil size. This discrepancy may be due to the inaccuracy of the used subjective reports as well as the imprecise report of timing and dosage of caffeine consumption in our study.

Our evidence against the occurrence of sex differences contrasts with the work of Harley & Sliney [47], who found that female subjects had significantly larger pupil sizes (0.30 mm on average) than male subjects under outdoor conditions. On the other hand, our results are in line with Winn et al. [28], who found no sex differences in their sample. The body of evidence for sex differences in pupil size therefore remains inconclusive. However, the lack of evidence in our data for iris colour as an influence factor for pupil size is consistent with both reports[28, 47].

### Further limitations

In addition to the list of study restrictions acknowledged in the Stage 1 Registered Report (see *Materials and Methods*), our study is limited in the following ways:

- *Time-resolution.* Given our temporal sampling resolution (10-second intervals), we may not have been able to capture some of the faster dynamics of pupil size, possibly stemming from cone signalling pathways [17, 18]. A finer time resolution would allow a more detailed analysis of the fast response aspects of pupil size regulation and the velocity of adaptation.
- *Exhaustive light scenarios.* We have not sampled all possible light scenarios and have limited our design to those that balance feasibility with full generalisability. As the actual statistical properties of the human spectral diet are unknown, this generalisability remains a construct.
- *Carry-over effects.* We acknowledged the omission of light history effects in Stage 1 and further found high autocorrelations values for pupil size and mEDI in our field data (see Suppl. Figure 4). It is thus plausible that our data include carry-over effects so that long-term pupil effects are mixed into the instantaneous measurements. However, this is a feature of naturalistic exposures in the real world. A clean but non-naturalistic design could employ counterbalanced sequences of light levels.

## Conclusion

While acknowledging the limitations and high data loss, this study is the first to the authors’ knowledge to investigate the pupillary light reflex under naturalistic light and task conditions enabled by a fully wearable measurement setup. Under these dynamic real-world conditions, the study sheds light on the factors influencing human pupil physiology in a sex-balanced, age-diverse sample. Using a Bayesian approach, we confirmed previous laboratory findings on senile miosis, the dose-response relationship between pupil size and light intensity, and the superiority of melanopsin-weighted measures over photopic illuminance for predicting steady-state pupil size. Our post-hoc analyses show moderate to strong evidence that iris colour, reported caffeine consumption, and sex did not substantially influence pupil size under these settings. Taken together, the data provide a strong case for considering age in personalised lighting solutions and for extending the use of melanopsin-weighted light measures to assess real-world lighting conditions for visual and non-visual functions.

## Table captions

**Table 1. *Participant characteristics***. Sample properties for participants that were included (n=83), excluded after the trial (n=23) and excluded before the trial (n=7). Age, sex, visual aid status and Body Mass Index (BMI) were based on self-report. Iris colour and weather during the trial were rated by the experimenter. The season was derived from the date of testing. Numerical variables are given as “*mean* (*standard deviation*)”, categorical variables are given as “*count* (*%*)”, missing values (“0 (NA%”)) result from participants being excluded before those items were surveyed.

**Table 2. *Confirmatory hypothesis tests (n=83, log_10_-transformed).*** The tested models are given in Wilkinson-Rogers’ notation (column 2), with the key variables coloured in red. The Likelihood ratios between the tested models are given as Bayes factors in scientific notation (column 3), along with the proportional error in percent (pe %; column 4), while the interpretation of the likelihood ratios follows the standard categorisations [58]. Here, the sample from the adjusted data loss threshold (75%, n=83) and log_10_-transformed light data from the field condition were used. For the initial data threshold with reduced sample (50%, n=63) and non-transformed light data, see results display in Suppl. tables 8-10. Abbreviations: CH = confirmatory hypothesis; 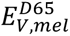 = melanopic Equivalent Daylight Illuminance; *E_V_* = photopic illuminance; BF_10_ = Bayes Factor comparing H1 with H0; % pe = proportional error in percent.

**Table 3. *Exploratory hypothesis tests (n=83, log_10_-transformed).*** The tested models are given in Wilkinson-Rogers’ notation (column 2), with the key variables coloured in red. The Likelihood ratios between the tested models are given as Bayes factors in scientific notation (column 3), along with the proportional error in percent (pe %; column 4), while the interpretation of the likelihood ratios follows the standard categorisations [58]. Here, the sample from the adjusted data loss threshold (75%, n=83) and log_10_-transformed light data from the field condition were used. Abbreviations: EH = exploratory hypothesis; 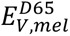 = melanopic Equivalent Daylight Illuminance; BF_10_ = Bayes Factor comparing H1 with H0; % pe = proportional error in percent.

## Authors’ Contributions

The authors made the following contributions. Rafael Robert Lazar: Conceptualization, Data curation, Formal Analysis, Investigation, Methodology, Project administration, Supervision, Visualization, Validation, Writing –original draft, Writing – review & editing; Josefine Degen: Investigation; Ann-Sophie Fiechter: Investigation; Aurora Monticelli: Investigation; Manuel Spitschan: Conceptualization, Data curation, Funding acquisition, Methodology, Project administration, Resources, Supervision, Visualization, Validation, Writing – original draft, Writing – review & editing.

## Acknowledgements

During parts of this work, M.S. was supported by a Sir Henry Wellcome Trust Fellowship (Wellcome Trust 204686/Z/16/Z) and a Junior Research Fellowship from Linacre College, University of Oxford. R.L. is funded by the European Training Network LIGHTCAP (project number 860613) under the Marie Skłodowska-Curie actions framework H2020-MSCA-ITN-2019. This study was funded by an investigator-initiated research project supported by Ocean Insight. We thank Dimitri Hawrylenko, Majlinda Maliqi, and Sebastian Saraceno for their assistance in the data collection, Dr Johannes Zauner for his helpful comments on the analysis code, and the anonymous reviewers for their comments in the Stage 1 submission.

## Footnotes

1 Due to difficulties in filling the older age groups during the COVID pandemic, we later opened recruitment to more than 16 per age group to achieve a large enough total sample. In total, n=113 people were invited to the health and eligibility screening of whom n=106 were assigned to participate in the experimental protocol (see Suppl. Figure 1).

2 No participants with IOLs were recruited in this study.

3 After Stage 1 IPA: we decided to adopt the more widely established melanopsin sensitivity-weighted light intensity unit melanopic Equivalent Daylight Illuminance (melanopic EDI or mEDI) instead of melanopic irradiance. This does not affect any of our test results, as the two measures are simply a linear transformation of one another (mEDI = melanopic irradiance ∗ 1.32621318911359).

4 After Stage 1 IPA: Our dataset revealed a higher proportion of pupil data exclusion than expected from the pilot runs. Thus, we decided to additionally use an adjusted threshold of ‘acceptable’ data loss proportion per participant to address high participant attrition. The second data loss threshold was determined in a data-driven approach (see *Results under Data quality checks, Proportion of excluded data*), resulting in a 75% threshold. For completeness and transparency, we report the results of our hypothesis tests in the reduced sample (n=63, 14354 valid observations, 50% threshold) and full sample (n=83, 17199 valid observations, 75% threshold).

5 Correction: An unrelated and wrongfully cited study by Yung, et al. (1990) in Stage 1 was replaced with the correct reference to Snel & Lorist (2011) in Stage 2.

6 Iris colour, biological sex, and caffeine consumption per body weight were included in the exploratory post-hoc analysis.

7 We followed the standard categorisation of Bayes factor strength of evidence according to Jeffreys (1961) [58] but omitted to note the interpretation for BF_10_>100 (“decisive evidence”).

8 During analysis (after Stage 1 IPA and data collection), we realised a peculiarity in our originally formulated linear mixed model analyses. When predicting pupil size with linear light intensity data (mEDI/melanopic irradiance and photopic illuminance), these variables violate the assumptions of linear regression models, specifically the assumptions for linearity, homogeneity of variance and collinearity (see Suppl. Fig. 5 B, C and E). However, when log10-transforming the light data before including them in the linear models, the assumptions approximately met (cf. Suppl. Fig. 6). Following these results, we log_10_-transformed our light data (more details see Linear Regression Assumptions under Results) For completeness and transparency, we report the results of our hypothesis tests with the log10-transformed and linear, non-transformed data. The two alternative methods produce qualitatively similar results, though BayesFactors are larger when employing the log10-transformation.

9 During analysis (after Stage 1 IPA and data collection), we realised a peculiarity in our originally formulated linear mixed model analyses. When predicting pupil size with linear light intensity data (mEDI/melanopic irradiance and photopic illuminance), these variables violate the assumptions of linear regression models, specifically the assumptions for linearity, homogeneity of variance and collinearity (see Suppl. Fig. 5 B, C and E). However, when log10-transforming the light data before including them in the linear models, the assumptions approximately met (cf. Suppl. Fig. 6). Following these results, we log_10_-transformed our light data (more details see Linear Regression Assumptions under Results) For completeness and transparency, we report the results of our hypothesis tests with the log10-transformed and linear, non-transformed data. The two alternative methods produce qualitatively similar results, though BayesFactors are larger when employing the log10-transformation.

10 During analysis (after Stage 1 IPA and data collection), we realised a peculiarity in our originally formulated linear mixed model analyses. When predicting pupil size with linear light intensity data (mEDI/melanopic irradiance and photopic illuminance), these variables violate the assumptions of linear regression models, specifically the assumptions for linearity, homogeneity of variance and collinearity (see Suppl. Fig. 5 B, C and E). However, when log10-transforming the light data before including them in the linear models, the assumptions approximately met (cf. Suppl. Fig. 6). Following these results, we log_10_-transformed our light data (more details see Linear Regression Assumptions under Results) For completeness and transparency, we report the results of our hypothesis tests with the log10-transformed and linear, non-transformed data. The two alternative methods produce qualitatively similar results, though BayesFactors are larger when employing the log10-transformation.

11 During analysis (after Stage 1 IPA and data collection), we realised a peculiarity in our originally formulated linear mixed model analyses. When predicting pupil size with linear light intensity data (mEDI/melanopic irradiance and photopic illuminance), these variables violate the assumptions of linear regression models, specifically the assumptions for linearity, homogeneity of variance and collinearity (see Suppl. Fig. 5 B, C and E). However, when log10-transforming the light data before including them in the linear models, the assumptions approximately met (cf. Suppl. Fig. 6). Following these results, we log_10_-transformed our light data (more details see Linear Regression Assumptions under Results) For completeness and transparency, we report the results of our hypothesis tests with the log10-transformed and linear, non-transformed data. The two alternative methods produce qualitatively similar results, though BayesFactors are larger when employing the log10-transformation.

12 During analysis (after Stage 1 IPA and data collection), we realised a peculiarity in our originally formulated linear mixed model analyses. When predicting pupil size with linear light intensity data (mEDI/melanopic irradiance and photopic illuminance), these variables violate the assumptions of linear regression models, specifically the assumptions for linearity, homogeneity of variance and collinearity (see Suppl. Fig. 5 B, C and E). However, when log10-transforming the light data before including them in the linear models, the assumptions approximately met (cf. Suppl. Fig. 6). Following these results, we log_10_-transformed our light data (more details see Linear Regression Assumptions under Results) For completeness and transparency, we report the results of our hypothesis tests with the log10-transformed and linear, non-transformed data. The two alternative methods produce qualitatively similar results, though BayesFactors are larger when employing the log10-transformation.

13 See footnote 4

