## Supplementary Material for "Regulation of pupil size in natural vision across the human lifespan"

### Supplementary Information

***Pre-processing.*** The data preparation was conducted with Unix Shell and Python scripts (2.7.12) that coordinated the processing of spectral data and estimation of pupil size, accessing "Pupil Core" software (Pupil Labs GmbH, 2019) in the process. The scripts were run on a Linux-Ubuntu (18.04.3 LTS) based high-performance desktop computer (32 GB RAM, Intel CoreTM i7-9700K CPU @3.60 HGZ × 8). During pre-processing, each sampling session – comprising a calibration video, pupil images and irradiance spectra – was processed separately after the respective experimentation. To use the pupil labs software, the still images of the pupil were pasted together with the calibration video via python. When the pre-processing script accessed the Pupil Player software to run the pupil size estimations from the assembled video, the settings had to be manually fine-tuned to the respective recordings. Firstly, an "area of interest" was set for each set of pupil video plus images to reduce misestimation of pupil size stemming from pixel disruptions outside of that area. Secondly, pupil size limits were configured, so that all pupil size values in the video were within the minimum and maximum values during all eye movements. Finally, the "intensity range", i.e., the minimum "blackness" of a pixel to be regarded as the pupil, was set. Here, it was critical to minimize the range to a value in which the pupil remained fully covered in detection while keeping erroneous pixel detections outside of the pupil as low as possible.
Once processed, each session was summarized in a CSV file, yielding pupil diameter, pupil detection confidence, melanopic irradiance and photopic illuminance values for every sample taken in 10-sec intervals. These csv-files formed the basis for the data analysis.

***Statistical software.*** Linear Mixed Model analyses with Bayes factors were conducted in R (version 4.3.1) [1], incorporating the lmbf function from the BayesFactor package (version 0.9.12-4.5) [2]. As specified in the Wilkinson-Rogers notation of the tested models in Stage 1, we classified "sex" and "id" as random factors. We then employed 10 000 iterations of Monte Carlo sampling to estimate each model's Bayes factor and then utilized the "compare" function to relate the full model with the null model to obtain the likelihood ratio ($BF_{10}$). All estimations were made reproducible by initializing the pseudorandom number generator at the same set point (Mersenne-Twister generator, using "set.seed(20230703)"). All figures generated in R used the "ggplot2" package (version 3.4.3) [3].

**Supplementary figures**

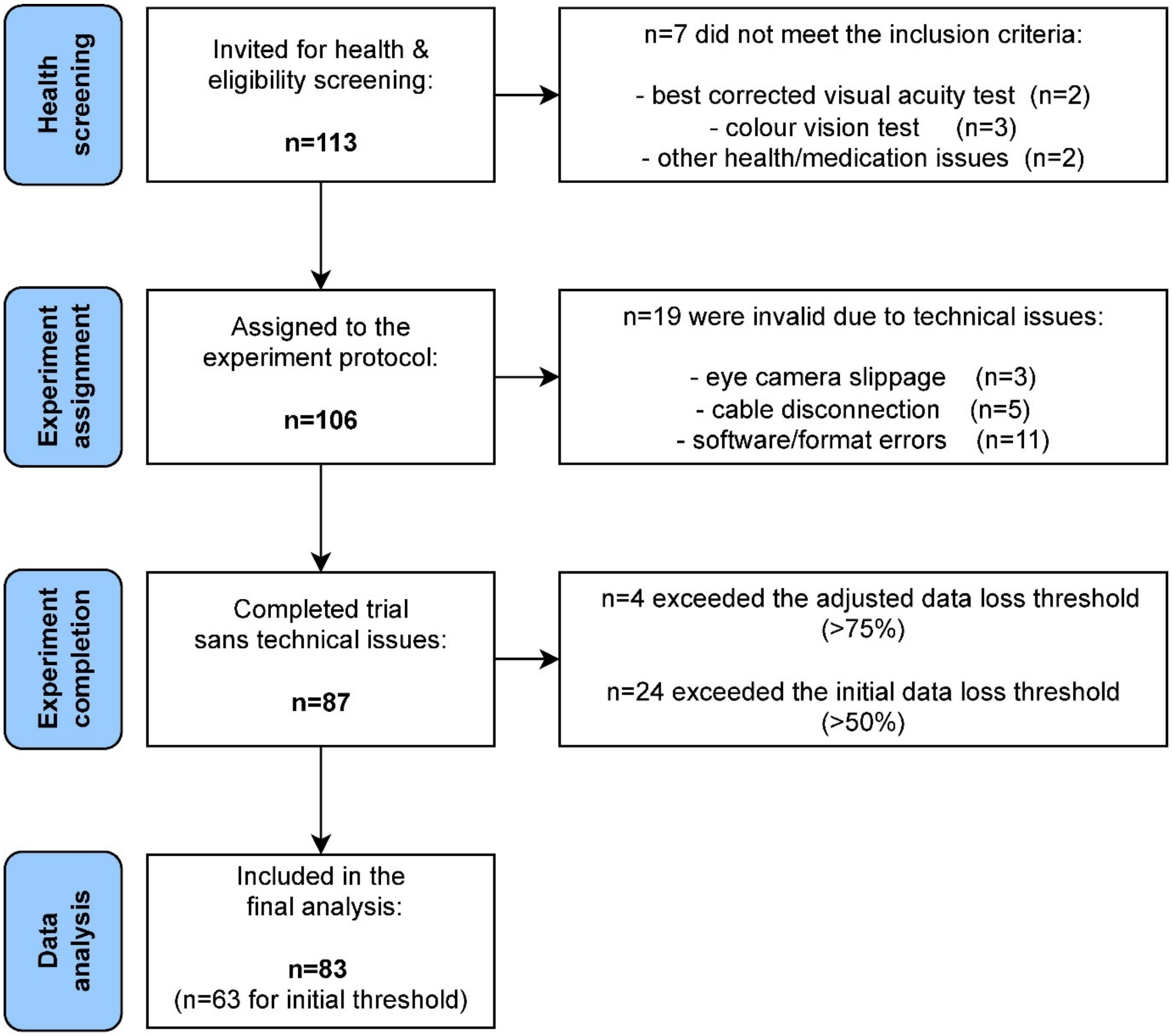


**Suppl. Figure 1**: Exclusion Flow diagram. Depiction of the recruitment process from the health screening to data analysis, including reasons for participant exclusion. The adjusted data loss threshold of 75% was applied, resulting in n=4 additional exclusions, retaining n=83 datasets. With the initial data loss threshold of 50%, n=24 additional participants are excluded, retaining 63 datasets.

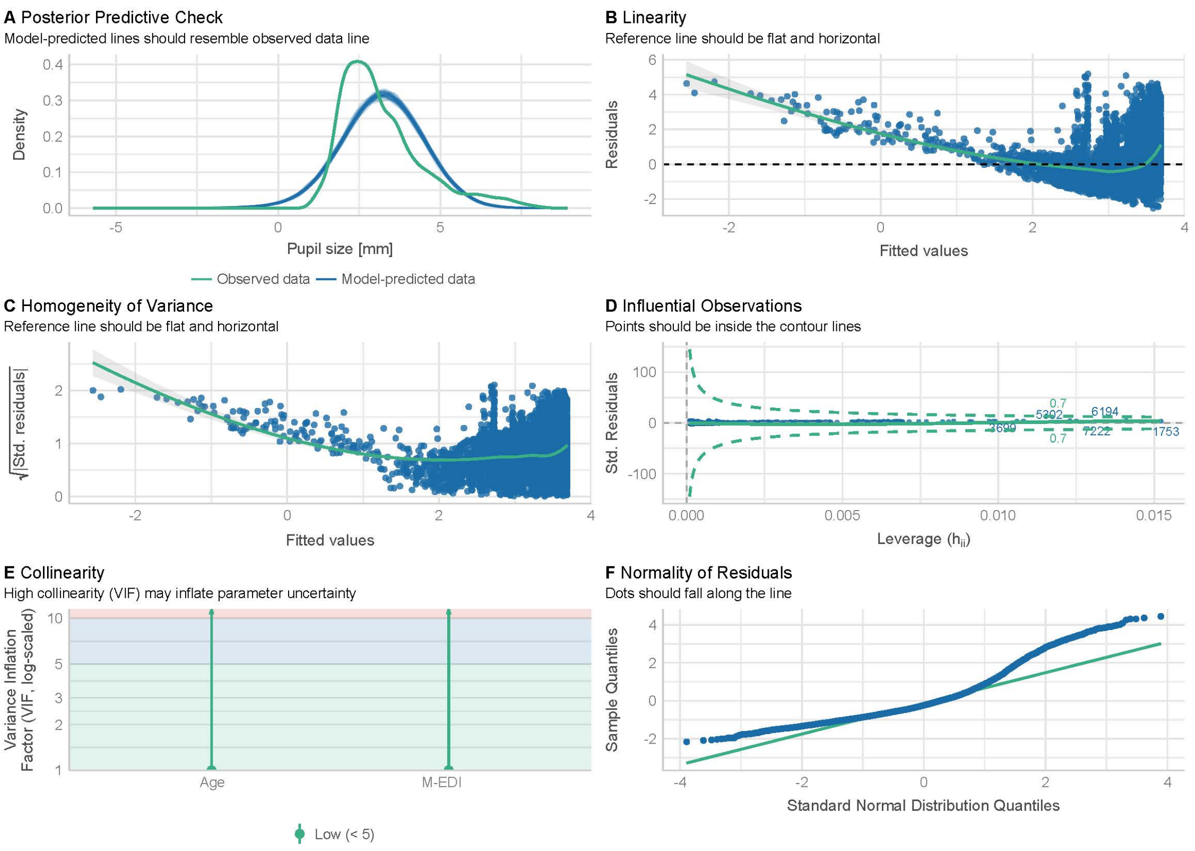


**Suppl. Figure 2**: (***A-E***) Visual check of linear regression assumptions – non-transformed light data (mEDI, n=83). ***A*** Posterior predictive check comparing the density of predicted pupil size data with observed data. ***B*** Linearity check, plotting residuals as a function of fitted values. ***C*** Homogeneity of variance (homoscedasticity) check, plotting the square root of standardized residuals as a function of fitted values. ***D*** Influential observations (outliers) check. ***E*** Collinearity (variance inflation) check. ***F*** Normality of Residuals check. Clear violations of the assumptions are visible in ***B***, ***C*** and ***E.***

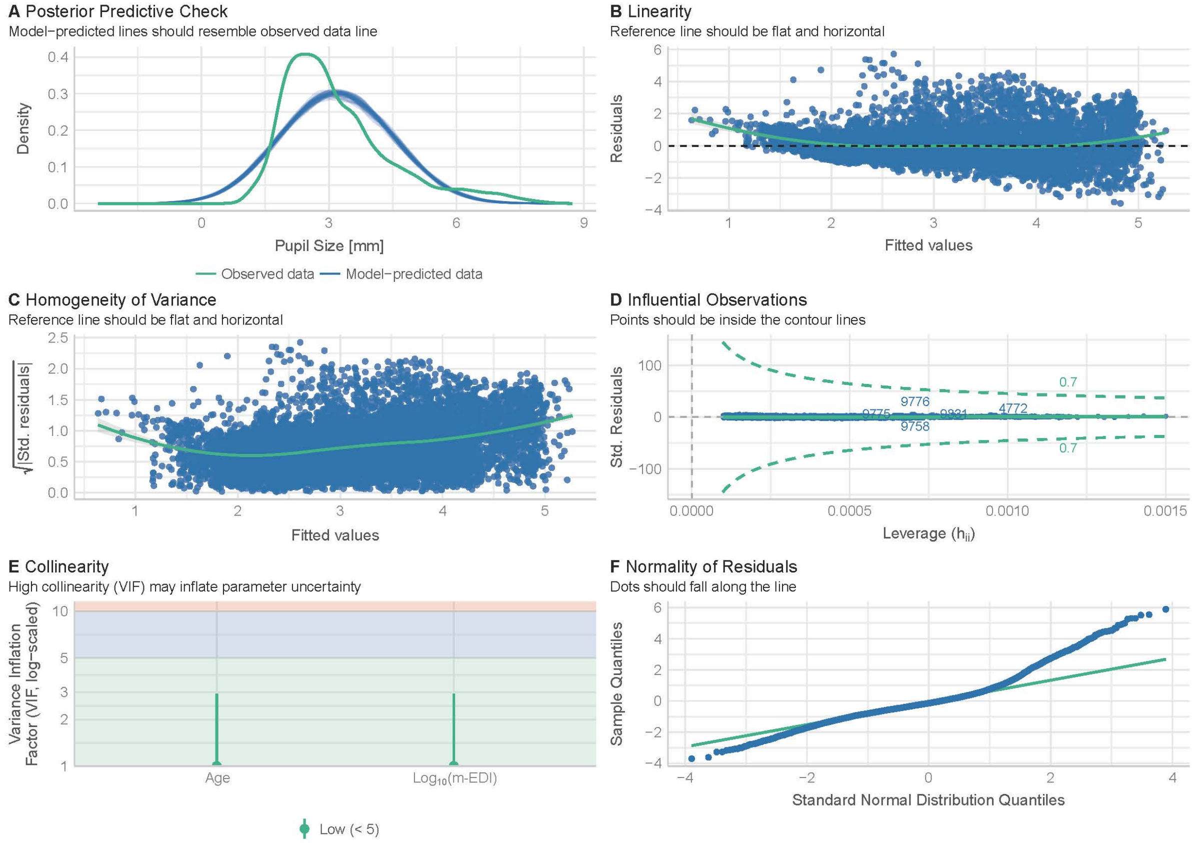


**Suppl. Figure 3**: (***A***-***E***) Visual check of linear regression assumptions – $log_{10}$-transformed light data ($log_{10}$(mEDI), n=83). ***A*** Posterior predictive check comparing the density of predicted pupil size data with observed data. ***B*** Linearity check, plotting residuals as a function of fitted values. ***C*** Homogeneity of variance (homoscedasticity) check, plotting the square root of standardized residuals as a function of fitted values. ***D*** Influential observations (outliers) check. ***E*** Collinearity (variance inflation) check. ***F*** Normality of Residuals check. The transformed data are approximately consistent with linear regression assumptions.

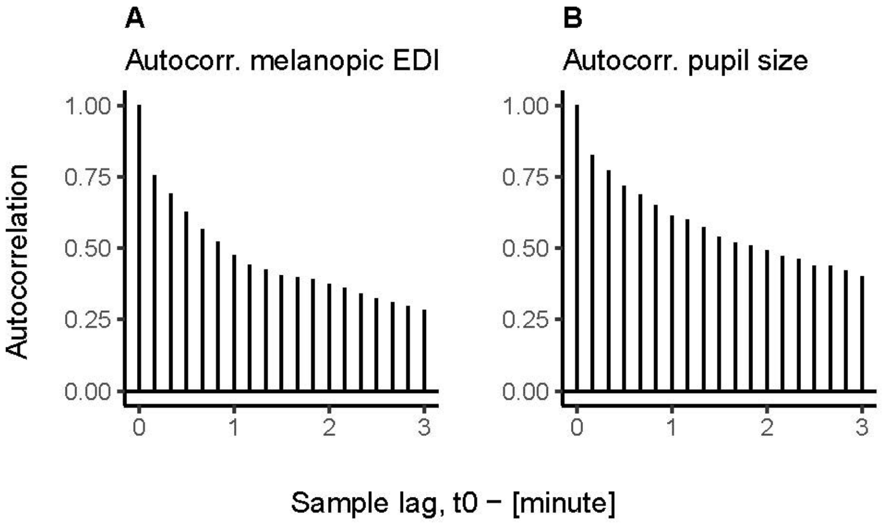


**Suppl. Figure 4**: (***A***-***B***) Autocorrelation of mEDI (***A)*** and pupil size (***B***) across a three-minute time lag, including all light conditions (n=83). Concurrent samples of pupil size and mEDI are highly autocorrelated with the sample 10 seconds prior (r>0.75) and remain slightly autocorrelated (r>0.25) with the sample three minutes before. Abbreviations: Autocorr. = autocorrelation, t0 = concurrent sample.

### Supplementary tables

**Suppl. Table 1.** ***Protocol.*** Tasks in the protocol (column 3) are specified regarding their light source ("a"=artificial, "n"=natural"; column 4) and location (column 5) across timestamps (columns 1 & 2).

| **Time stamp** | **Time elapsed h:mm** | **Task specification** | **Light source (artificial / natural)** | **Location** |
|---|---|---|---|---|
| - | 0:00 | Set-up and calibration | **a** | Laboratory |
| | **0:00** | **Start sampling** | **a** | Laboratory |
| 1. | 0:00 | Baseline – 10-min dark adaptation | **-** | Laboratory |
| 2. | 0:10 | Quality check laboratory condition:<br>4 spectrally different light phases in 3 intensities (=12 settings) | **a** | Laboratory |
| 3. | 0:25 | Navigation to office room | **a** | Corridor |
| 4. | 0:26 | Indoor tasks: laptop tasks, questionnaires, text reading & window gazing | **a+n** | Office |
| 5. | 0:34 | Additional indoor tasks: navigating & hand washing | **a+n** | Office, Corridor & bathroom |
| 6. | 0:40 | Navigation to outdoors | **a+n** | Corridor |
| 7. | 0:41 | Outdoor tasks: walking, conversing, text reading, inspecting environment. | **n** | Outdoor on campus |
| 8. | 0:58 | Navigation back to laboratory | **a +n** | Corridor |
| 9. | **1:00** | **Stop sampling** | **a** | Laboratory |
| | | Removing devices and end of experiment | **a** | Laboratory |

**Suppl. Table 2**. ***Additional participant characteristics.*** Further sample properties for participants that were included (n=83), excluded after the trial (n=23) and excluded before the trial (n=7). All variables but "trial start time" (noted by the experimenter) and "time between wake-up and trial" were assessed via self-report (see *Recorded demographic and ancillary information.* Numerical variables are given as "*mean* (*standard deviation*)", categorical variables are given as "*count* (*%*)", and missing values ("NA (NA)") result from participants being excluded before those items were surveyed.

| **Variable** | **Included**, N = 83[1] | **Excluded after trial**, N = 23[1] | **Excluded before trial**, N = 7[1] |
|---|---|---|---|
| **Trial start time [h]** | 12.25 (2.35) | 12.20 (2.52) | 11.59 (1.51) |
| **Handedness** | | | |
| Right | 75 (90%) | 17 (74%) | 6 (86%) |
| Left | 7 (8.4%) | 3 (13%) | 1 (14%) |
| Both | 1 (1.2%) | 3 (13%) | 0 (0%) |
| **Midpoint of sleep (corrected for sleep compensation) [h]** | 2.36 (3.27) | 3.11 (2.88) | NA (NA) |
| **Time between wake-up and trial [h]** | 5.00 (2.43) | 4.42 (3.19) | NA (NA) |
| **Sleep duration before trial [h]** | 7.92 (2.48) | 8.63 (4.16) | NA (NA) |
| **Average daily sleep duration [h]** | 8.28 (2.37) | 8.52 (2.21) | NA (NA) |
| **Estimated acute caffeine consumption [mg/prior** | 66.55 (80.59) | 68.41 (62.99) | NA (NA) |
| **Estimated habitual caffeine consumption [mg/day]** | 174.76 (172.64) | 237.64 (204.49) | NA (NA) |
| **Subjective sleepiness mid-trial [KSS scale, 1-9]** | 3.73 (1.87) | 3.91 (2.07) | NA (NA) |
| **Subjective sleepiness post-trial [KSS scale, 1-9]** | 3.19 (1.59) | 3.50 (2.26) | NA (NA) |

[1] Mean (SD); n (%)

**Suppl. Table 3**. ***Pupil size data summary per participant (n=83).*** Data loss values are computed as the ratio between excluded observations (due to invalid pupil or light data) and all observations based on the while dataset (all conditions). Summarized pupil data is separated between data from the field conditions (left) and the laboratory plus dark adaptation conditions used for positive control (right). Here, only observations with valid pupil data were used. Abbreviations: lab = laboratory; dark adapt. = dark adaptation; min.= minimum; med.= median; max. = maximum.

| | | | Field data | | | Positive control data (lab. & dark-adapt.) | | |
|---|---|---|---|---|---|---|---|---|
| ID | Age | Data loss ratio [%] | Min. pupil size [mm] | Med. pupil size [mm] | Max. pupil size [mm] | Min. pupil size [mm] | Med. pupil size [mm] | Max. pupil size [mm] |
| SP001 | 28 | 25.00 | 2.04 | 2.67 | 7.95 | 2.61 | 4.78 | 9.00 |
| SP003 | 22 | 39.60 | 1.53 | 2.78 | 6.27 | 1.83 | 5.34 | 6.64 |
| SP004 | 25 | 55.78 | 2.08 | 2.98 | 7.18 | 2.28 | 6.46 | 8.14 |
| SP005 | 19 | 40.99 | 1.23 | 2.02 | 7.26 | 1.96 | 3.31 | 7.76 |
| SP006 | 30 | 47.13 | 1.24 | 2.31 | 7.25 | 2.18 | 4.76 | 7.87 |
| SP008 | 25 | 17.05 | 2.14 | 3.13 | 8.03 | 2.47 | 7.58 | 8.39 |
| SP009 | 25 | 23.51 | 1.23 | 2.20 | 7.00 | 2.04 | 5.58 | 7.17 |
| SP010 | 26 | 46.53 | 1.57 | 2.68 | 7.88 | 2.81 | 7.68 | 8.00 |
| SP011 | 25 | 66.00 | 1.43 | 3.59 | 7.46 | 2.29 | 3.63 | 7.46 |
| SP012 | 33 | 46.17 | 1.81 | 2.66 | 7.90 | 2.47 | 5.07 | 8.14 |
| SP013 | 34 | 47.18 | 1.63 | 2.15 | 6.94 | 2.07 | 4.55 | 7.31 |
| SP014 | 22 | 42.12 | 1.03 | 2.73 | 7.73 | 1.99 | 3.65 | 8.98 |
| SP015 | 26 | 35.19 | 1.55 | 2.16 | 4.99 | 1.59 | 3.63 | 5.62 |
| SP017 | 45 | 69.12 | 2.56 | 2.93 | 5.03 | 1.80 | 3.14 | 5.24 |
| SP018 | 32 | 29.05 | 1.52 | 2.47 | 6.29 | 1.70 | 3.72 | 6.58 |
| SP020 | 22 | 48.35 | 1.47 | 2.49 | 6.41 | 2.22 | 3.50 | 7.34 |
| SP021 | 40 | 29.74 | 1.62 | 3.90 | 5.65 | 2.71 | 4.76 | 6.30 |
| SP022 | 26 | 37.57 | 1.32 | 2.42 | 5.90 | 1.83 | 4.74 | 7.12 |
| SP024 | 39 | 30.65 | 1.06 | 2.26 | 4.78 | 1.52 | 2.95 | 4.91 |
| SP025 | 25 | 45.67 | 1.58 | 3.25 | 8.22 | 2.26 | 4.34 | 8.06 |
| SP026 | 24 | 26.20 | 2.57 | 3.84 | 7.73 | 2.29 | 5.27 | 7.96 |
| SP027 | 24 | 25.57 | 2.34 | 3.35 | 7.16 | 2.53 | 6.80 | 7.63 |
| SP028 | 48 | 56.71 | 1.67 | 2.02 | 4.06 | 2.10 | 2.77 | 6.03 |
| SP029 | 24 | 43.55 | 1.30 | 2.03 | 5.15 | 1.88 | 4.44 | 6.12 |
| SP030 | 18 | 32.58 | 2.45 | 3.81 | 6.56 | 2.36 | 4.99 | 7.23 |
| SP031 | 38 | 23.95 | 2.31 | 3.08 | 7.69 | 2.32 | 6.15 | 7.89 |
| SP032 | 22 | 41.34 | 1.25 | 3.08 | 6.59 | 1.45 | 7.18 | 7.61 |
| SP033 | 57 | 18.42 | 1.29 | 1.76 | 5.21 | 1.67 | 4.21 | 5.86 |
| SP034 | 51 | 25.98 | 1.79 | 2.49 | 5.50 | 1.95 | 5.12 | 6.51 |
| SP037 | 36 | 50.00 | 1.87 | 3.58 | 8.32 | 2.26 | 5.13 | 8.98 |
| SP038 | 18 | 34.59 | 1.94 | 3.55 | 7.73 | 2.35 | 5.46 | 8.28 |
| SP039 | 27 | 60.82 | 2.24 | 4.13 | 8.03 | 1.41 | 5.36 | 8.54 |
| SP041 | 21 | 44.89 | 2.36 | 4.60 | 7.50 | 2.63 | 7.24 | 8.02 |

| | | | | | | | | |
|---|---|---|---|---|---|---|---|---|
| SP043 | 21 | 34.91 | 2.07 | 3.19 | 6.58 | 2.34 | 3.74 | 7.98 |
| SP044 | 36 | 23.74 | 1.85 | 3.06 | 6.43 | 2.04 | 4.50 | 6.64 |
| SP046 | 22 | 44.33 | 1.81 | 4.28 | 6.70 | 2.33 | 6.62 | 7.31 |
| SP047 | 20 | 64.02 | 1.21 | 3.54 | 6.74 | 1.27 | 6.50 | 6.71 |
| SP048 | 20 | 35.64 | 1.91 | 3.82 | 8.26 | 1.65 | 4.49 | 8.84 |
| SP049 | 46 | 30.23 | 1.03 | 1.88 | 3.42 | 1.24 | 2.20 | 4.25 |
| SP050 | 85 | 47.93 | 1.87 | 3.13 | 6.79 | 2.08 | 3.47 | 6.92 |
| SP053 | 36 | 48.84 | 1.99 | 3.35 | 5.79 | 1.91 | 3.39 | 6.69 |

| | | | **Field data** | | | **Positive control data (lab. & dark adapt.)** | | |
|---|---|---|---|---|---|---|---|---|
| **ID** | **Age** | **Data loss ratio [%]** | **Min. pupil size [mm]** | **Med. pupil size [mm]** | **Max. pupil size [mm]** | **Min. pupil size [mm]** | **Med. pupil size [mm]** | **Max. pupil size [mm]** |
| SP054 | 55 | 42.21 | 1.99 | 3.05 | 5.98 | 2.34 | 3.46 | 7.13 |
| SP055 | 39 | 52.85 | 1.30 | 2.65 | 6.59 | 2.08 | 6.89 | 7.18 |
| SP056 | 60 | 22.13 | 1.21 | 2.34 | 5.41 | 1.70 | 4.62 | 5.63 |
| SP057 | 54 | 70.64 | 1.25 | 2.30 | 5.47 | 2.20 | 6.74 | 7.23 |
| SP059 | 44 | 40.59 | 1.34 | 3.35 | 6.14 | 1.35 | 3.29 | 6.83 |
| SP063 | 61 | 64.30 | 1.29 | 2.54 | 3.96 | 3.57 | 4.74 | 5.05 |
| SP064 | 80 | 54.20 | 1.38 | 2.58 | 4.79 | 2.01 | 2.84 | 5.28 |
| SP065 | 38 | 37.75 | 1.57 | 2.87 | 7.34 | 2.03 | 3.24 | 7.59 |
| SP066 | 48 | 55.36 | 1.46 | 2.56 | 5.64 | 1.30 | 3.28 | 6.01 |
| SP067 | 57 | 25.75 | 1.13 | 2.42 | 3.61 | 1.40 | 3.09 | 4.38 |
| SP068 | 40 | 33.87 | 1.04 | 3.42 | 6.67 | 1.39 | 6.26 | 7.33 |
| SP070 | 54 | 30.84 | 1.54 | 2.59 | 6.02 | 1.99 | 5.45 | 6.47 |
| SP071 | 21 | 32.09 | 1.84 | 4.31 | 8.23 | 2.78 | 6.30 | 8.42 |
| SP072 | 62 | 15.83 | 2.21 | 3.36 | 7.01 | 2.44 | 5.16 | 7.24 |
| SP073 | 61 | 60.00 | 1.19 | 2.70 | 5.34 | 1.25 | 3.26 | 6.67 |
| SP074 | 64 | 10.82 | 1.35 | 2.72 | 4.00 | 1.48 | 3.27 | 3.88 |
| SP076 | 87 | 58.01 | 1.65 | 2.52 | 4.70 | 1.90 | 2.87 | 4.98 |
| SP077 | 38 | 26.44 | 2.46 | 3.59 | 6.44 | 2.08 | 5.52 | 6.96 |
| SP080 | 41 | 26.74 | 1.79 | 2.46 | 5.53 | 1.63 | 4.90 | 5.89 |
| SP081 | 20 | 57.68 | 2.02 | 3.04 | 7.72 | 2.32 | 7.44 | 7.94 |
| SP082 | 23 | 61.14 | 1.97 | 3.10 | 7.42 | 2.12 | 3.87 | 7.63 |
| SP083 | 20 | 26.24 | 2.26 | 3.44 | 7.95 | 2.34 | 5.05 | 8.02 |
| SP084 | 21 | 40.39 | 2.58 | 3.42 | 7.65 | 1.02 | 3.97 | 8.30 |
| SP085 | 26 | 36.77 | 1.79 | 3.33 | 6.41 | 1.71 | 6.79 | 8.10 |
| SP087 | 45 | 56.22 | 2.12 | 3.05 | 5.79 | 1.43 | 3.26 | 5.85 |
| SP089 | 22 | 41.71 | 1.59 | 3.25 | 7.22 | 1.20 | 3.78 | 7.86 |
| SP090 | 24 | 54.95 | 1.06 | 3.58 | 6.77 | 1.33 | 3.90 | 7.35 |
| SP091 | 19 | 35.73 | 1.89 | 4.33 | 7.39 | 2.13 | 5.21 | 7.59 |
| SP092 | 24 | 36.16 | 1.67 | 2.37 | 7.16 | 1.56 | 4.49 | 8.06 |
| SP093 | 19 | 45.04 | 1.66 | 2.64 | 5.05 | 1.07 | 3.28 | 5.96 |
| SP095 | 26 | 52.83 | 1.48 | 2.35 | 6.67 | 2.18 | 3.64 | 7.86 |
| SP096 | 18 | 44.72 | 1.67 | 3.14 | 7.24 | 2.68 | 6.34 | 8.46 |

| | | | | | | | | |
|---|---|---|---|---|---|---|---|---|
| SP097 | 20 | 46.56 | 1.16 | 3.92 | 8.11 | 2.70 | 6.36 | 8.56 |
| SP098 | 24 | 37.29 | 1.42 | 2.66 | 6.11 | 1.87 | 3.79 | 6.73 |
| SP100 | 21 | 41.89 | 1.35 | 2.20 | 5.13 | 2.12 | 3.59 | 6.36 |
| SP101 | 34 | 52.80 | 1.81 | 3.40 | 5.26 | 2.12 | 6.83 | 7.32 |
| SP108 | 18 | 27.70 | 1.61 | 2.80 | 7.42 | 2.37 | 5.03 | 8.16 |
| SP109 | 54 | 41.33 | 1.80 | 3.27 | 5.67 | 1.52 | 4.89 | 6.16 |
| SP110 | 42 | 60.76 | 2.15 | 2.91 | 5.34 | 1.95 | 2.59 | 6.22 |
| SP111 | 32 | 35.56 | 1.61 | 2.09 | 4.89 | 1.80 | 5.13 | 5.28 |
| SP112 | 77 | 25.26 | 2.04 | 2.94 | 7.91 | 2.86 | 6.75 | 8.33 |
| SP113 | 67 | 19.72 | 1.51 | 2.18 | 4.21 | 1.02 | 2.98 | 5.00 |

**Suppl. Table 4**. ***Light data summary per participant (n=83)***. Summarized mEDI data is separated between data from the field conditions (left) and the laboratory and dark adaptation conditions used for positive control (right). Only observations with valid pupil and light data were used. Abbreviations: lab = laboratory; dark adapt. = dark adaptation; mEDI = melanopic Equivalent Daylight Illuminance; min.= minimum; med.= median; max. = maximum.

| | | Field data | | | Positive control data (lab. & dark adapt.) | | |
|---|---|---|---|---|---|---|---|
| **ID** | **Age** | **Min. mEDI [lx]** | **Med. mEDI [lx]** | **Max. mEDI [lx]** | **Min. mEDI [lx]** | **Med. mEDI [lx]** | **Max. mEDI [lx]** |
| SP001 | 28 | 1 | 1798 | 37697 | ~0 | 65 | 1956 |
| SP003 | 22 | 2 | 584 | 32244 | ~0 | 62 | 1922 |
| SP004 | 25 | 3 | 1032 | 29374 | ~0 | 63 | 1848 |
| SP005 | 19 | 3 | 2207 | 29490 | ~0 | 57 | 1728 |
| SP006 | 30 | 2 | 2258 | 11347 | ~0 | 59 | 824 |
| SP008 | 25 | 1 | 939 | 9440 | ~0 | 68 | 2079 |
| SP009 | 25 | 2 | 1798 | 15867 | ~0 | 61 | 1850 |
| SP010 | 26 | 4 | 2528 | 27431 | ~0 | 8 | 1878 |
| SP011 | 25 | 2 | 87 | 7237 | ~0 | 13 | 1713 |
| SP012 | 33 | 4 | 1678 | 24324 | ~0 | 14 | 1829 |
| SP013 | 34 | 2 | 2197 | 9695 | ~0 | 61 | 1865 |
| SP014 | 22 | 3 | 641 | 4996 | ~0 | 50 | 1669 |
| SP015 | 26 | 4 | 708 | 9110 | ~0 | 55 | 1779 |
| SP017 | 45 | 2 | 37 | 188 | ~0 | 62 | 1872 |
| SP018 | 32 | 2 | 546 | 44912 | ~0 | 60 | 1830 |
| SP020 | 22 | 1 | 1997 | 12965 | ~0 | 54 | 1722 |
| SP021 | 40 | 2 | 164 | 4153 | ~0 | 56 | 1674 |
| SP022 | 26 | 3 | 341 | 19136 | ~0 | 54 | 1701 |
| SP024 | 39 | 2 | 279 | 35081 | ~0 | 59 | 1871 |
| SP025 | 25 | 2 | 65 | 13734 | ~0 | 57 | 1752 |
| SP026 | 24 | 3 | 145 | 1015 | ~0 | 54 | 1636 |
| SP027 | 24 | 3 | 47 | 557 | ~0 | 35 | 744 |
| SP028 | 48 | 3 | 1059 | 22307 | ~0 | 61 | 1824 |
| SP029 | 24 | 2 | 1200 | 28463 | ~0 | 537 | 1829 |
| SP030 | 18 | 1 | 103 | 2805 | ~0 | 57 | 1689 |
| SP031 | 38 | 2 | 205 | 2450 | ~0 | 61 | 1797 |
| SP032 | 22 | 3 | 438 | 11216 | ~0 | 59 | 795 |
| SP033 | 57 | 3 | 796 | 22017 | ~0 | 64 | 1905 |
| SP034 | 51 | 3 | 566 | 17421 | ~0 | 61 | 1855 |
| SP037 | 36 | 2 | 214 | 27528 | ~0 | 63 | 1891 |
| SP038 | 18 | 4 | 178 | 37967 | ~0 | 51 | 1647 |
| SP039 | 27 | 2 | 68 | 15583 | ~0 | 56 | 1748 |
| SP041 | 21 | 3 | 50 | 36436 | ~0 | 58 | 1795 |
| SP043 | 21 | 3 | 256 | 5652 | ~0 | 64 | 1857 |
| SP044 | 36 | 3 | 85 | 3292 | ~0 | 57 | 1777 |
| SP046 | 22 | 3 | 34 | 30663 | ~0 | 56 | 1698 |

| | | | | | | | |
|---|---|---|---|---|---|---|---|
| SP047 | 20 | 1 | 92 | 13221 | ~0 | 15 | 702 |
| SP048 | 20 | 2 | 181 | 3707 | ~0 | 54 | 1593 |
| SP049 | 46 | 3 | 234 | 30214 | ~0 | 60 | 1796 |
| SP050 | 85 | 3 | 168 | 28320 | ~0 | 62 | 1913 |
| SP053 | 36 | 3 | 54 | 11292 | ~0 | 57 | 1617 |
| SP054 | 55 | 3 | 81 | 28853 | ~0 | 56 | 1680 |

| | | Field data | | | Positive control data (lab. & dark adapt.) | | |
|---|---|---|---|---|---|---|---|
| **ID** | **Age** | **Min. mEDI [lx]** | **Med. mEDI [lx]** | **Max. mEDI [lx]** | **Min. mEDI [lx]** | **Med. mEDI [lx]** | **Max. mEDI [lx]** |
| SP055 | 39 | 5 | 374 | 19249 | ~0 | 64 | 826 |
| SP056 | 60 | 3 | 294 | 16712 | ~0 | 61 | 1851 |
| SP057 | 54 | 4 | 1250 | 40197 | ~0 | 42 | 1819 |
| SP059 | 44 | 3 | 55 | 5675 | ~0 | 62 | 1889 |
| SP063 | 61 | 3 | 48 | 23058 | ~0 | NA | -Inf |
| SP064 | 80 | 2 | 169 | 32407 | ~0 | 60 | 1849 |
| SP065 | 38 | 3 | 101 | 12692 | ~0 | 118 | 1776 |
| SP066 | 48 | 3 | 80 | 8369 | ~0 | 59 | 1828 |
| SP067 | 57 | 3 | 43 | 29198 | ~0 | 57 | 1760 |
| SP068 | 40 | 3 | 38 | 4734 | ~0 | 59 | 1801 |
| SP070 | 54 | 3 | 358 | 42979 | ~0 | 59 | 1740 |
| SP071 | 21 | 3 | 96 | 5778 | ~0 | 49 | 1449 |
| SP072 | 62 | 3 | 70 | 4389 | ~0 | 57 | 1698 |
| SP073 | 61 | 2 | 25 | 1832 | ~0 | 39 | 1269 |
| SP074 | 64 | 2 | 31 | 5451 | ~0 | 60 | 1832 |
| SP076 | 87 | 2 | 601 | 27967 | ~0 | 59 | 1782 |
| SP077 | 38 | 3 | 112 | 1261 | ~0 | 63 | 1861 |
| SP080 | 41 | 3 | 161 | 4092 | ~0 | 63 | 1911 |
| SP081 | 20 | 3 | 1173 | 40780 | ~0 | 14 | 1600 |
| SP082 | 23 | 3 | 583 | 2215 | ~0 | 54 | 1685 |
| SP083 | 20 | 1 | 436 | 3490 | ~0 | 54 | 1632 |
| SP084 | 21 | 2 | 196 | 2375 | ~0 | 63 | 1721 |
| SP085 | 26 | 3 | 88 | 6611 | ~0 | 61 | 1845 |
| SP087 | 45 | 2 | 23 | 2390 | ~0 | 57 | 1649 |
| SP089 | 22 | 2 | 411 | 4692 | ~0 | 62 | 1780 |
| SP090 | 24 | 2 | 112 | 12305 | ~0 | 47 | 1672 |
| SP091 | 19 | 1 | 216 | 1135 | ~0 | 57 | 1767 |
| SP092 | 24 | 3 | 727 | 24002 | ~0 | 59 | 1816 |
| SP093 | 19 | 2 | 192 | 4567 | ~0 | 60 | 1890 |
| SP095 | 26 | 3 | 201 | 16052 | ~0 | 505 | 1807 |
| SP096 | 18 | 3 | 145 | 3877 | ~0 | 18 | 1763 |
| SP097 | 20 | 2 | 122 | 26897 | ~0 | 59 | 1848 |
| SP098 | 24 | 3 | 268 | 10682 | ~0 | 58 | 1777 |
| SP100 | 21 | 1 | 845 | 33649 | ~0 | 59 | 1777 |

| | | | | | | | |
|---|---|---|---|---|---|---|---|
| SP101 | 34 | 3 | 35 | 34985 | ~0 | 34 | 1699 |
| SP108 | 18 | 2 | 505 | 21362 | ~0 | 55 | 1635 |
| SP109 | 54 | 3 | 18 | 483 | ~0 | 56 | 1687 |
| SP110 | 42 | 3 | 312 | 915 | ~0 | 62 | 1800 |
| SP111 | 32 | 3 | 391 | 4326 | ~0 | 674 | 1717 |
| SP112 | 77 | 3 | 908 | 28101 | ~0 | 60 | 1838 |
| SP113 | 67 | 3 | 104 | 3660 | ~0 | 60 | 1866 |

**Suppl. Table 5.** ***Log$_{10}$(mEDI) transformation test (n=83).*** The tested models are given in Wilkinson-Rogers' notation (column 2), with the key variables coloured in red. The Likelihood ratios between the tested models are given as Bayes factors in scientific notation (column 3), along with the proportional error in per cent (pe %; column 4), while the interpretation of the likelihood ratios follows the standard categorisations [4]. The test was calculated in the field data with the adjusted data threshold (75%, n=83). Abbreviations: $E_{V,mel}^{D65}$ = melanopic Equivalent Daylight Illuminance; $E_V$ = photopic illuminance; BF$_{10}$ = Bayes Factor comparing H1 with H0; % pe = proportional error in percent.

| Test | H$_1$/H$_0$ Model comparison [Wilkinson-Rogers' notation] | BF$_{10}$ | % pe | Interpretation |
|---|---|---|---|---|
| Log$_{10}$(m-EDI) Vs. m-EDI | $\frac{\text{Pupil size} = \log_{10}(E_{V,mel}^{D65}) + Age + (1\|Id) + (1\|Sex)}{\text{Pupil size} = E_{V,mel}^{D65} + Age + (1\|Id) + (1\|Sex)}$ | 5.678918e+996 | ±3.55% | decisive evidence for the log$_{10}$ model |

**Suppl. Table 6.** ***Positive control tests (n=83).*** The tested models are given in Wilkinson-Rogers' notation (column 2), with the key variables coloured in red. The Likelihood ratios between the tested models are given as Bayes factors in scientific notation (column 3), along with the proportional error in per cent (pe %; column 4), while the interpretation of the likelihood ratios follows the standard categorisations [4]. Positive control tests for CH1 and CH2 were calculated in the laboratory dataset with the adjusted data threshold (75%, n=83; rows 1-4), including $\log_{10}$-transformed (rows 1-2) as well as linear (non-transformed) light data (rows 3-4). The positive control test for CH3 was computed in the dark adaptation dataset with the adjusted data loss threshold (75%, n=83; row 5). Abbreviations: CH = Confirmatory Hypothesis; lab = laboratory data; dark = dark-adaptation data; $E_{V,mel}^{D65}$ = melanopic Equivalent Daylight Illuminance; $E_V$ = photopic illuminance; $BF_{10}$ = Bayes Factor comparing H1 with H0; % pe = proportional error in percent.

| Positive control tests | $H_1/H_0$ Model comparison [Wilkinson-Rogers' notation] | $BF_{10}$ | % pe | Interpretation |
|---|---|---|---|---|
| ***CH1 – Lab log₁₀*** | $\frac{\text{Pupil size} = \log_{10}(E_{V,mel}^{D65}) + Age + (1\|Id) + (1\|Sex)}{\text{Pupil size} = Age + (1\|Id) + (1\|Sex)}$ | 1.090096e+127 | ±2.57% | decisive evidence for CH1 vs. null model |
| **CH2 – Lab log₁₀** | $\frac{\text{Pupil size} = \log_{10}(E_{V,mel}^{D65}) + Age + (1\|Id) + (1\|Sex)}{\text{Pupil size} = \log_{10}(E_V)\ Age + (1\|Id) + (1\|Sex)}$ | 4.524662e+24 | ±2.13% | decisive evidence for CH2 vs. null model |
| **CH1 – Lab linear** | $\frac{\text{Pupil size} = E_{V,mel}^{D65} + Age + (1\|Id) + (1\|Sex)}{\text{Pupil size} = Age + (1\|Id) + (1\|Sex)}$ | 1.553889e+170 | ±2.55% | decisive evidence for CH1 vs. null model |
| **CH2 – Lab linear** | $\frac{\text{Pupil size} = E_{V,mel}^{D65} + Age + (1\|Id) + (1\|Sex)}{\text{Pupil size} = E_V + Age + (1\|Id) + (1\|Sex)}$ | 3.366674e+99 | ±1.85% | decisive evidence for CH2 vs. null model |
| **CH3 – Dark** | $\frac{\text{Pupil size} = Age + (1\|Id) + (1\|Sex)}{\text{Pupil size} = (1\|Id) + (1\|Sex)}$ | 208318.5 | ±1.44% | decisive evidence for CH3 vs. null model |

**Suppl. Table 7.** ***Positive control tests (n=63).*** The tested models are given in Wilkinson-Rogers' notation (column 2), with the key variables coloured in red. The Likelihood ratios between the tested models are given as Bayes factors in scientific notation (column 3), along with the proportional error in per cent (pe %; column 4), while the interpretation of the likelihood ratios follows the standard categorisations [4]. Positive control tests for CH1 and CH2 were calculated in the laboratory dataset with the initial data threshold (50%, n=63; rows 1-4), including $\log_{10}$-transformed (rows 1-2) as well as linear (non-transformed) light data (rows 3-4). The positive control test for CH3 was computed in the dark adaptation dataset with the inital data loss threshold (50%, n=63; row 5). Abbreviations: CH = Confirmatory Hypothesis; lab = laboratory data; dark = dark-adaptation data; $E_{V,mel}^{D65}$ = melanopic Equivalent Daylight Illuminance; $E_V$ = photopic illuminance; $BF_{10}$ = Bayes Factor comparing H1 with H0; % pe = proportional error in percent.

| Hypo-thesis | $H_1/H_0$ Model comparison [Wilkinson-Rogers' notation] | $BF_{10}$ | % pe | Interpretation |
|---|---|---|---|---|
| ***CH1 – Lab log10*** | $\frac{\text{Pupil size} = \log_{10}(E_{V,mel}^{D65}) + Age + (1\|Id) + (1\|Sex)}{\text{Pupil size} = Age + (1\|Id) + (1\|Sex)}$ | 2.801948e+116 | ±2.06% | decisive evidence for CH1 vs. null model |
| **CH2 – Lab log10** | $\frac{\text{Pupil size} = \log_{10}(E_{V,mel}^{D65}) + Age + (1\|Id) + (1\|Sex)}{\text{Pupil size} = \log_{10}(E_V)\ Age + (1\|Id) + (1\|Sex)}$ | 4.262853e+27 | ±2.31% | decisive evidence for CH2 vs. null model |
| ***CH1 – Lab* linear** | $\frac{\text{Pupil size} = E_{V,mel}^{D65} + Age + (1\|Id) + (1\|Sex)}{\text{Pupil size} = Age + (1\|Id) + (1\|Sex)}$ | 1.136029e+153 | ±2.05% | decisive evidence for CH1 vs. null model |
| **CH2 – Lab linear** | $\frac{\text{Pupil size} = E_{V,mel}^{D65} + Age + (1\|Id) + (1\|Sex)}{\text{Pupil size} = E_V + Age + (1\|Id) + (1\|Sex)}$ | 6.961333e+91 | ±1.82% | decisive evidence for CH2 vs. null model |
| **CH3 – Dark** | $\frac{\text{Pupil size} = Age + (1\|Id) + (1\|Sex)}{\text{Pupil size} = (1\|Id) + (1\|Sex)}$ | 192.7163 | ±1.36% | decisive evidence for CH3 vs. null model |

**Suppl. Table 8.** ***Confirmatory hypothesis tests (n=83, linear light data).*** The tested models are given in Wilkinson-Rogers' notation (column 2), with the key variables coloured in red. The Likelihood ratios between the tested models are given as Bayes factors in scientific notation (column 3), along with the proportional error in percent (pe %; column 4), while the interpretation of the likelihood ratios follows the standard categorisations [4]. Here, the sample from the adjusted data loss threshold (75%, n=83) and linear (non-transformed) light data from the field condition were used. Abbreviations: CH = confirmatory hypothesis; $E_{V,mel}^{D65}$ = melanopic Equivalent Daylight Illuminance; $E_V$ = photopic illuminance; $BF_{10}$ = Bayes Factor comparing H1 with H0; % pe = proportional error in percent.

| Hypo-thesis | $H_1/H_0$ Model comparison [Wilkinson-Rogers' notation] | $BF_{10}$ | % pe | Interpretation |
|---|---|---|---|---|
| **CH1** | $\frac{\text{Pupil size} = E_{V,mel}^{D65} + Age + (1\|Id) + (1\|Sex)}{\text{Pupil size} = Age + (1\|Id) + (1\|Sex)}$ | 2.642936e+291 | ±2.78% | decisive evidence for CH1 vs. null model |
| **CH2** | $\frac{\text{Pupil size} = E_{V,mel}^{D65} + Age + (1\|Id) + (1\|Sex)}{\text{Pupil size} = E_V + Age + (1\|Id) + (1\|Sex)}$ | 10708989894 | ±3.91% | decisive evidence for CH2 vs. null model |
| **CH3** | $\frac{\text{Pupil size} = E_{V,mel}^{D65} + Age + (1\|Id) + (1\|Sex)}{\text{Pupil size} = E_{V,mel}^{D65} + (1\|Id) + (1\|Sex)}$ | 2221.5 | ±2.66% | decisive evidence for CH3 vs. null model |

**Suppl. Table 9.** ***Confirmatory hypothesis tests (n=63, $log_{10}$-transformed light data).*** The tested models are given in Wilkinson-Rogers' notation (column 2), with the key variables coloured in red. The Likelihood ratios between the tested models are given as Bayes factors in scientific notation (column 3), along with the proportional error in per cent (pe %; column 4), while the interpretation of the likelihood ratios follows the standard categorisations [4]. Here, the sample from the initial data loss threshold (50%, n=63) and $\log_{10}$-transformed light data from the field condition were used. Abbreviations: CH = confirmatory hypothesis; $E_{V,mel}^{D65}$ = melanopic Equivalent Daylight Illuminance; $E_V$ = photopic illuminance; $BF_{10}$ = Bayes Factor comparing H1 with H0; % pe = proportional error in percent.

| Hypo-thesis | $H_1/H_0$ Model comparison[1] [Wilkinson-Rogers' notation] | $BF_{10}$ | % pe | Interpretation |
|---|---|---|---|---|
| **CH1** | $\frac{\text{Pupil size} = \log_{10}(E_{V,mel}^{D65}) + Age + (1\|Id) + (1\|Sex)}{\text{Pupil size} = Age + (1\|Id) + (1\|Sex)}$ | 2.404706e+1076 | ±2.24% | decisive evidence for CH1 vs. null model |
| **CH2** | $\frac{\text{Pupil size} = \log_{10}(E_{V,mel}^{D65}) + Age + (1\|Id) + (1\|Sex)}{\text{Pupil size} = \log_{10}(E_V)\ Age + (1\|Id) + (1\|Sex)}$ | 1.782022e+19 | ±2.06% | decisive evidence for CH2 vs. null model |
| **CH3** | $\frac{\text{Pupil size} = \log_{10}(E_{V,mel}^{D65}) + Age + (1\|Id) + (1\|Sex)}{\text{Pupil size} = \log_{10}(E_{V,mel}^{D65}) + (1\|Id) + (1\|Sex)}$ | 124.465 | ±2.12% | decisive evidence for CH3 vs. null model |

**Suppl. Table 10.** ***Confirmatory hypothesis tests (n=63, linear light data).*** The tested models are given in Wilkinson-Rogers' notation (column 2), with the key variables coloured in red. The Likelihood ratios between the tested models are given as Bayes factors in scientific notation (column 3), along with the proportional error in per cent (pe %; column 4), while the interpretation of the likelihood ratios follows the standard categorisations [4]. Here, the sample from the initial data loss threshold (50%, n=63) and linear (non-transformed) light data from the field condition were used. Abbreviations: CH = confirmatory hypothesis; $E_{V,mel}^{D65}$ = melanopic Equivalent Daylight Illuminance; $E_V$ = photopic illuminance; $BF_{10}$ = Bayes Factor comparing H1 with H0; % pe = proportional error in percent.

| Hypo-thesis | $H_1/H_0$ Model comparison [Wilkinson-Rogers' notation] | $BF_{10}$ | % pe | Interpretation |
|---|---|---|---|---|
| **CH1** | $\frac{\text{Pupil size} = E_{V,mel}^{D65} + Age + (1\|Id) + (1\|Sex)}{\text{Pupil size} = Age + (1\|Id) + (1\|Sex)}$ | 1.031709e+242 | ±2.24% | decisive evidence for CH1 vs. null model |
| **CH2** | $\frac{\text{Pupil size} = E_{V,mel}^{D65} + Age + (1\|Id) + (1\|Sex)}{\text{Pupil size} = E_V + Age + (1\|Id) + (1\|Sex)}$ | 1921990 | ±1.93% | decisive evidence for CH2 vs. null model |
| **CH3** | $\frac{\text{Pupil size} = E_{V,mel}^{D65} + Age + (1\|Id) + (1\|Sex)}{\text{Pupil size} = E_{V,mel}^{D65} + (1\|Id) + (1\|Sex)}$ | 13.992 | ±1.89% | strong evidence for CH3 vs. null model |
