## Supplementary material for "Regulation of pupil size in natural vision across the human lifespan": Table 1

| Variable | **Included**, N = 83[1] | **Excluded after trial**, N = 23[1] | **Excluded before trial**, N = 7[1] |
|---|---|---|---|
| **Age [y]** | 35.70 (17.16) | 37.87 (20.13) | 52.00 (21.54) |
| **Age group [y]** | | | |
| 18-24 | 30 (36%) | 9 (39%) | 0 (0%) |
| 25-34 | 18 (22%) | 5 (22%) | 2 (29%) |
| 35-44 | 13 (16%) | 0 (0%) | 1 (14%) |
| 45-54 | 9 (11%) | 3 (13%) | 0 (0%) |
| 55-64 | 8 (9.6%) | 5 (22%) | 1 (14%) |
| >64 | 5 (6.0%) | 1 (4.3%) | 3 (43%) |
| **Sex** | | | |
| Female | 43 (52%) | 17 (74%) | 3 (43%) |
| Male | 40 (48%) | 6 (26%) | 4 (57%) |
| **Uses visual aid** | | | |
| No | 57 (69%) | 16 (70%) | 3 (43%) |
| Yes - myopia correction | 18 (22%) | 5 (22%) | 2 (29%) |
| Yes - hyperopia correction | 8 (9.6%) | 2 (8.7%) | 2 (29%) |
| **Wearing contact lenses during trial** | 16 (19%) | 5 (22%) | 2 (29%) |
| **BMI** | 22.96 (3.47) | 22.07 (3.93) | 26.39 (5.79) |
| **Iris colour** | | | |
| Blue | 25 (30%) | 9 (39%) | 0 (NA%) |
| Hazel/Green | 17 (20%) | 8 (35%) | 0 (NA%) |
| Brown | 41 (49%) | 6 (26%) | 0 (NA%) |
| **Weather during trial** | | | |
| Light rain | 9 (11%) | 0 (0%) | 0 (NA%) |
| Very cloudy | 16 (19%) | 7 (30%) | 0 (NA%) |
| Cloudy | 14 (17%) | 1 (4.3%) | 0 (NA%) |
| Somewhat cloudy | 11 (13%) | 5 (22%) | 0 (NA%) |
| Sunny | 33 (40%) | 10 (43%) | 0 (NA%) |
| **Season** | | | |
| Spring | 7 (8.4%) | 5 (22%) | 1 (14%) |
| Summer | 40 (48%) | 9 (39%) | 3 (43%) |
| Autumn | 16 (19%) | 5 (22%) | 0 (0%) |
| Winter | 20 (24%) | 4 (17%) | 3 (43%) |

[1] Mean (SD); n (%)
