## Supplementary material for "Regulation of pupil size in natural vision across the human lifespan": Table 2

| Hypo-  thesis | H_1_/H_0_ Model comparison  [Wilkinson-Rogers’ notation] | BF_10_ | % pe | Interpretation |
| --- | --- | --- | --- | --- |
| **CH1** | $\frac{Pupil size=\log_{10} (E_{V,mel}^{D65})+ Age +(1\vert Id) +(1\vert Sex)}{Pupil size= Age +(1\vert Id) +(1\vert Sex)}$ | 1.543255e+1288 | ±2.2% | decisive evidence for CH1 vs. null model |
| **CH2** | $\frac{Pupil size=\log_{10} \left( E_{V,mel}^{D65} \right)+ Age +\left( 1 \vert Id \right)+\left( 1 \vert Sex \right)}{Pupil size=\log_{10} \left( E_{V} \right) Age +\left( 1 \vert Id \right)+\left( 1 \vert Sex \right)}$ | 1.458261e+21 | ±2.13% | decisive evidence for CH2 vs. null model |
| **CH3** | $\frac{Pupil size=\log_{10} \left( E_{V,mel}^{D65} \right)+ Age +\left( 1 \vert Id \right)+\left( 1 \vert Sex \right)}{Pupil size=\log_{10} \left( E_{V,mel}^{D65} \right)+\left( 1 \vert Id \right)+\left( 1 \vert Sex \right)}$ | 46076.190 | ±2.17% | decisive evidence for CH3 vs. null model |

**Table 2. *Confirmatory hypothesis tests*** ***(n=83, log_10_-transformed).***
