## Supplementary material for "Regulation of pupil size in natural vision across the human lifespan": Table 3

| Hypo-  thesis | H_1_/H_0_ Model comparison  [Wilkinson-Rogers’ notation] | BF_10_ | % pe | Interpretation |
| --- | --- | --- | --- | --- |
| **EH1** | $\frac{Pupil size=\log_{10} (E_{V,mel}^{D65})+ Age +(1\vert Id) +(1\vert Sex)}{Pupil size=\log_{10} (E_{V,mel}^{D65})+ Age +(1\vert Id)}$ | 0.070 | ±1.44% | strong evidence for the null model vs. EH1 |
| **EH2** | $\frac{Pupil size=\log_{10} \left( E_{V,mel}^{D65} \right)+ Age+\left( 1 \vert Id \right)+\left( 1 \vert Sex \right)+ (1\vert Iris colour)}{Pupil size=\log_{10} \left( E_{V,mel}^{D65} \right)+ Age+\left( 1 \vert Id \right)+\left( 1 \vert Sex \right)}$ | 0.017 | ±2.86% | strong evidence for the null model  vs. EH2 |
| **EH3** | $\frac{Pupil size=\log_{10} \left( E_{V,mel}^{D65} \right)+ Age ++\left( 1 \vert Id \right)+\left( 1 \vert Sex \right)+(Habitual caffeine/kg)}{Pupil size=\log_{10} \left( E_{V,mel}^{D65} \right)+ Age ++\left( 1 \vert Id \right)+\left( 1 \vert Sex \right)}$ | 0.131 | ±2.10% | moderate evidence for the null model  vs. EH3 |
| **EH4** | $\frac{Pupil size=\log_{10} \left( E_{V,mel}^{D65} \right)+ Age ++\left( 1 \vert Id \right)+\left( 1 \vert Sex \right)+(Acute caffeine/kg)}{Pupil size=\log_{10} \left( E_{V,mel}^{D65} \right)+ Age ++\left( 1 \vert Id \right)+\left( 1 \vert Sex \right)}$ | 0.176 | ±2.10% | moderate evidence for the null model  vs. EH4 |

**Table 3. *Exploratory hypothesis tests (n=83, log_10_-transformed).***
